# Odor-evoked Increases in Olfactory Bulb Mitral Cell Spiking Variability

**DOI:** 10.1101/2021.03.05.434131

**Authors:** Cheng Ly, Andrea K. Barreiro, Shree Hari Gautam, Woodrow L. Shew

**Author notes:** Correspondences should be addressed to Cheng Ly.

## Abstract

At the onset of sensory stimulation, the variability and co-variability of spiking activity is widely reported to decrease, especially in cortex. Considering the potential benefits of such decreased variability for coding, it has been suggested that this could be a general principle governing all sensory systems. Here we show that this is not so. We recorded mitral cells in olfactory bulb (OB) of anesthetized rats and found increased variability and co-variability of spiking at the onset of olfactory stimulation. Using models and analysis, we predicted that these increases arise due to network interactions within OB, without increasing variability of input from the nose. We tested and confirmed this prediction in awake animals with direct optogenetic stimulation of OB to circumvent the pathway through the nose. Our results establish increases in spiking variability at stimulus onset as a viable alternative coding strategy to the more commonly observed decreases in variability in many cortical systems.

**Summary:** The spiking variability of neural networks has important implications for how information is encoded to higher brain regions. It has been well documented by numerous labs in many cortical and motor regions that spiking variability decreases with stimulus onset, yet whether this principle holds in the olfactory bulb has not been tested. In stark contrast to this common view, we demonstrate that the onset of sensory input can cause an increase in the variability of neural activity in the mammalian olfactory bulb. We show this in both anesthetized and awake rodents. Furthermore, we use computational models to describe the mechanisms of this phenomenon. Our finding establish sensory evoked increases in spiking variability as a viable alternative.

## Introduction

The variability of spiking in response to sensory input can significantly impact the coding efficacy of downstream neurons. Numerous experimental observations in diverse sensory cortical regions have established that the onset of sensory input causes a drop in the average spiking variability (Churchland et al., 2010). Similarly, the spiking co-variability among simultaneously recorded pairs of neurons decreases in the evoked state (Doiron et al., 2016). There have been many theoretical studies detailing how neuron populations can have both increases in firing rate and decreases in (co-)variability at stimulus onset. The most prominent mechanistic explanations are succinctly summarized in Doiron et al. (2016), and include changes in noise levels via (pre-)synaptic transmission (Churchland et al., 2010; Ponce-Alvarez et al., 2013; Rosenbaum et al., 2013), altering transfer function of inputs to outputs (de la Rocha et al., 2007; Barreiro et al., 2010, 2012), and increases in correlated inhibition leading to cancellation of correlated inputs (Renart et al., 2010; Litwin-Kumar et al., 2011; Middleton et al., 2012; Tetzlaff et al., 2012; Ly et al., 2012).

Does stimulus-induced reduction of variability and co-variability generalize across all sensory neural systems? Here we first sought to answer this question in the olfactory bulb (**OB**), which processes sensory signals generated by olfactory receptor neurons (**ORN**) in the nose and passes this sensory information on to piriform cortex. Notably, the OB was not discussed in Churchland et al. (2010) nor in any of the experiments cited in Doiron et al. (2016) despite its crucial role in olfaction. We recorded single unit activity from anesthetized rats during precisely controlled olfactory stimulation. We were surprised to find that stimulus onset caused an increase in variability and co-variability of spiking among mitral cells (**MC**s) in OB.

Stimulus-induced increases in spiking variability and co-variability are not widely reported and are understudied (but see Kazama and Wilson (2009); Wanner and Friedrich (2020)). Moreover, the mechanisms governing spiking variability and co-variability in OB are not well understood. Thus, our next goals were to elucidate mechanisms of variability in the OB circuit, and more specifically, to clarify why the OB variability increases rather than decreases. We focus on two possible sources of variability. First, the input to OB is generated by olfactory receptor neurons, thus variability of ORN activity is one likely source of OB variability. It has been observed that ORN activity increases with stimulus intensity (Scott et al., 2007; Furudono et al., 2013), but the extent to which the variability (across trials or across glomeruli) of ORN input changes at stimulus onset is unclear. In *Drosophila*, changes in ORN inputs that occur with stimulus onset modulate spiking of projection neurons, the analog of MCs (Kazama and Wilson, 2009). A second important factor in spiking variability of recurrent networks, in general, is inhibition. Thus, we also explored the role of granule cells (**GC**, an important inhibitory neuron in OB) in OB variability. GC activity is known to affect spiking variability and co-variability in OB (Galán et al., 2006; Giridhar et al., 2011).

To this end, we developed and analyzed computational models of the OB. We use a relatively simple firing rate model that contains many known biophysical details of the OB, including having 3 important cell types (MC, GC, and periglomerular cells (**PGC**)), sub-network architecture, membrane time-constants, and synaptic dynamics (based on the Cleland OB models (Li and Cleland, 2013, 2017)). Our model enables a detailed computational approach for determining coupling strengths and ORN inputs that best match our data. We found for a given ORN variance regime that uncorrelated GC inhibition to MC must be strong, excitation to GC is weaker, and lateral inhibition providing shared input to multiple MCs must be weak. When accounting for three key spike statistics — time-varying firing rate, spike count variance and covariance — we found that the total error between model and data is minimized when the ORN input noise is fixed across spontaneous and evoked states. Thus our model predicts that ORN input noise does not need to increase in the evoked state to observe evoked increases in OB spiking variability.

We tested this prediction using previous experiments in awake mice (Bolding and Franks, 2018a,b). In these experiments, the optogenetic stimulation of OB acted as a surrogate input to OB, replacing ORN input, but with no increase in ORN variability at stimulus onset. In agreement with our model prediction, OB activity exhibited an increase in firing variability at the onset of optogenetic stimulation.

Taken together, experimental data and our modeling results establish increased variability at stimulus onset as a viable strategy for sensory systems and demonstrate how this occurs in OB.

## Results

We performed *in vivo* multi-electrode array recordings of the OB in the mitral cell layer of anesthetized rats (see **STAR Methods and Methods**) to capture both spontaneous and odor-evoked spiking activity of populations of putative MCs. This yielded a large number of cells and simultaneously recorded pairs of cells with heterogeneous trial-averaged spiking activity. Figure 1 summarizes the trial-averaged spiking with a food-like odor (Ethyl Butyrate), showing the population firing rate (peri-stimulus time histogram, **PSTH** in Fig.1**A**), spike count variance (Fig.1**B**) and spike count covariance (Fig.1**C**) over 100 ms time windows. The choice of 100 ms time window is an intermediate value between the shorter time scales (membrane time constants, AMPA, GABA_A_, etc.) and longer ones (NMDA, calcium and other ionic currents) relevant to the circuit. Details about the trial-averaged spiking statistics on a cell-by-cell, or pair-by-pair basis are shown in Figure 1**D–F**, where the time-averaged spike count is shown in each state (spontaneous and evoked).

**Figure 1:**
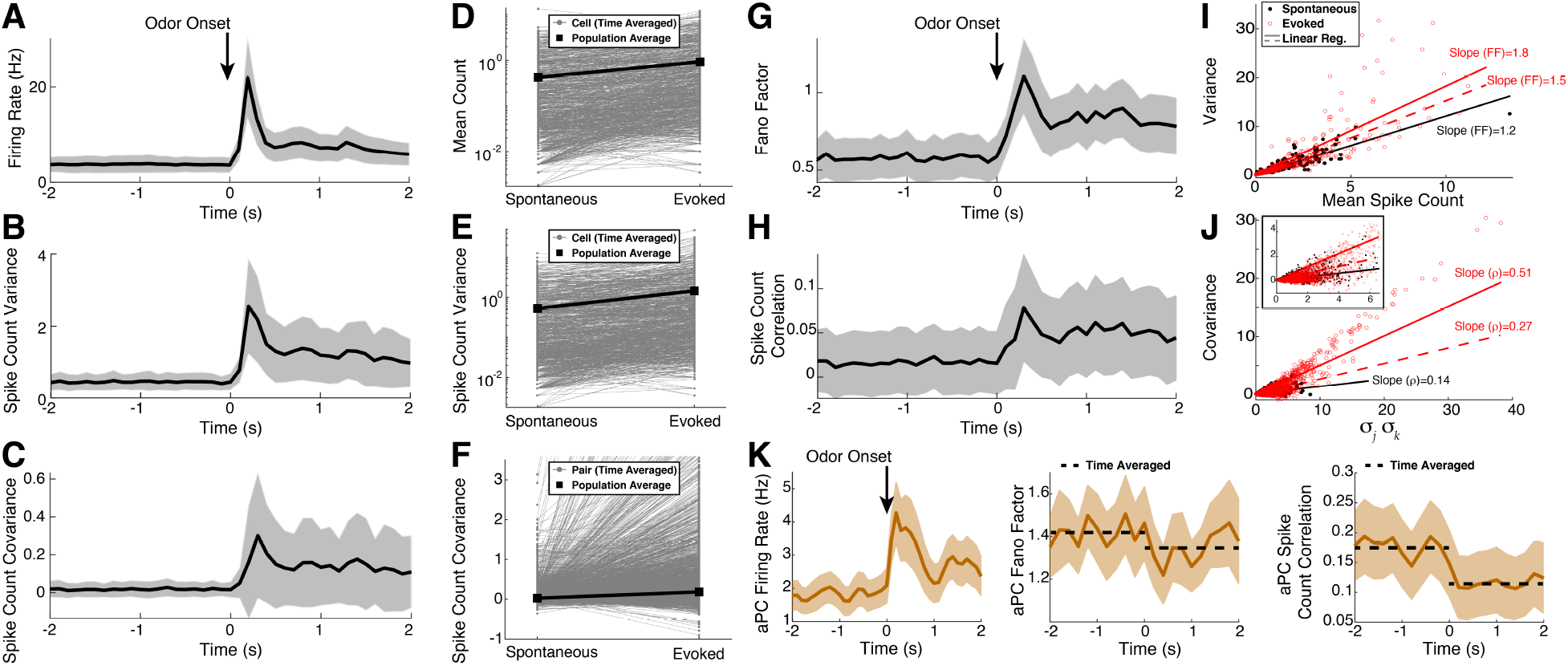
Spiking statistics from experiments. **A)–C)** Spiking statistics in spontaneous (*t* < 0) and odor-evoked states (*t* > 0); the firing rate (**A**), spike count variance (**B**) and covariance (**C**). **D)–F)** Trial- and time-averaged spike count statistics of individual cells/pairs in gray (black squares = population average) also generally increase. The mean firing rate (**D**), 918 out of 1120 cells, 82%), variance (**E**), 908 out of 1120 cells, 81.1%, vertical log-scale), and covariance (**F**) 10194 out of 17674 cell pairs, 57.7%) increase with odor. **G)–J) Scaled measures of variability shown for completeness**. Fano factor (**G**) and Pearson’s Correlation (**H**) of spike count increase with odor (Ethyl Butyrate). **I**) Spike count variance versus mean in spontaneous (black) and evoked (red) states, time-averaged in each state. Lines are regression of best least squares fit to data with 0 intercept, another way to measure Fano factor. Two red lines for evoked state: the solid line (slope of 1.83, *R*^2^ = 0.660) is regression for all data, lower line (red dashed, slope of 1.52, *R*^2^ = 0.657) is regression on a subset of data (evoked mean count ≤ largest spontaneous mean count) to account for larger mean counts in the evoked state (see mean-matching in Churchland et al. (2010)). Spontaneous Fano factor (slope) is 1.21 (*R*^2^ = 0.870) is smaller than evoked Fano factor. **J**) Spike count covariance versus product of standard deviations of spike counts. The slope of the regression line is another measure of Pearson’s correlation ρ. Spontaneous ρ = 0.14 (*R*^2^ = 0.385) and evoked ρ = 0.51 (*R*^2^ = 0.750); evoked with cut-off: ρ = 0.27 (*R*^2^ = 0.565). **K**) Spike statistics from simultaneously recording in the anterior piriform cortex (**aPC**) are consistent with other data: evoked decreases in Fano factor (Churchland et al., 2010; Miura et al., 2012) and correlation (Miura et al., 2012). Spike counts were binned in 100 ms half-overlapping windows, except for **K** where 200 ms was used. Data averaged over 10 trials and over cell population (1120 cells in **A, B, D, E, G**; 17674 pairs in **C, F, H**; 194 cells and 7395 pairs in **K**). Shaded gray/tan regions show relative population heterogeneity: μ ± 0.2std (standard deviation across the population/pairs after trial-averaging; 0.2 scale for visualization).

Consistent with many sensory and motor neurons, the odor-evoked firing rates are larger than the spontaneous firing rates on average. However, the spike count variability (population-averaged variance, Fig. 1**B**, and Fano factor, defined as variance over mean, Fig. 1**G**) *increases* in the odor-evoked state, in contrast to many cortical regions where spiking variability decreases with stimuli (Churchland et al., 2010). Similarly, we see on average an *increase* in spike count co-variability (population-averaged covariance, Fig. 1C, and Pearson’s correlation, Fig. 1**H**) in the evoked state compared to the spontaneous state, again in contrast to many cortical regions (Doiron et al., 2016). Another way to measure population Fano factor is via the slope of regression line (with intercept at the origin) for a cell’s variance against mean spike count; similarly correlation can also be measured by slope of regression line of covariance of pairs against product of standard deviations.

By these measures we again see evoked increases of these statistics (Fig. 1**I, J**). Churchland et al. (2010) accounted for the increase in mean activity in the evoked state to attribute a decrease in Fano factor to the spike count variance by a ‘mean-matching method’. Here we account for increases in mean spike count in a more crude and conservative way by performing linear regression for a subset of data, removing all cells/pairs that have evoked mean count larger than the largest spontaneous mean count and regressing on the resulting subset: we still observe evoked increases in Fano factor and correlation (red dotted lines in Fig. 1**I, J**). Although these scaled measures of variability increase with stimuli (see Fig. 1**G–J**), we focus on capturing the three unscaled statistics (PSTH, variance and covariance of spike counts) because they each directly effect common measures of neural coding (e.g. the Fisher information) whereas it is less clear how scaled statistics such as Fano factor and correlation relate to neural coding (Kohn et al., 2016).

Finally, Figure 1**K** shows that simultaneously recorded cells from anterior piriform cortex (**aPC**) are consistent with previous reports. Our data shows odor-evoked increases in the population firing rate (left), and odor-evoked decreases in Fano factor (middle, Churchland et al. (2010); Miura et al. (2012)) and correlation (right, Miura et al. (2012)).

### Summary of Modeling Results

Here we list the key results of our data-driven modeling, from sections **Firing rate model** to **Application to other experimental data**, because the technical details can be distracting.

- We initially consider 3 ways to vary the ORN input noise (Fig 2B) in our model to determine which best captures our data (PSTH, spike variance, spike covariance).
- We systematically vary coupling strengths between MCs and GCs and find similar coupling strength constraints for all ORN input noise regime (Fig 3D, Table 1, Fig 6E, Table 3).
- We find fixed ORN input noise captures our data best (Fig 3C). We probe other ORN input noise regimes to find this result is robust (Fig. 4A). Even with awake data, we find ORN input noise does not need to increase for evoked increases in MC spike variability (Fig 6D).

**Figure 2:**
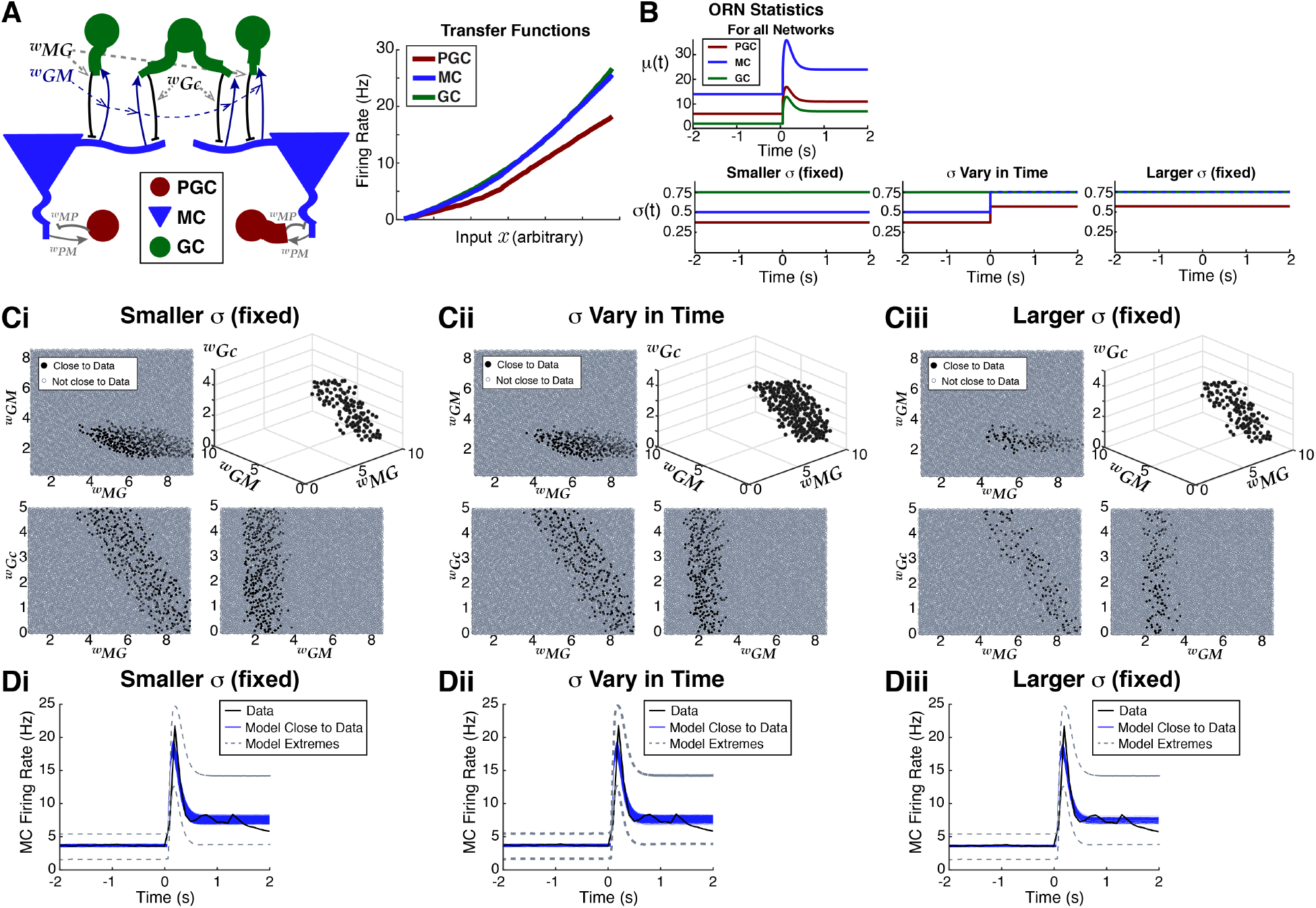
Feasible region in parameter space to match firing rate (PSTH) from data. **A**) Left: sub-network of 7 cells highlighting the 3 coupling strengths we systematically vary: *w*_*MG*_, *w*_*GM*_, and *w*_*Gc*_. **A**) Right: transfer functions used in our model, obtained numerically from Cleland OB models (Li and Cleland, 2013, 2017) (see **Supplementary Material: Details of the transfer function calculation** Figure S4C). **B**) Model of odor input from ORN (see Eq. (4)), having time-varying mean μ(t) (top) and 3 ways to set variance of ORN inputs (bottom): i) fixed but small *σ* (**Smaller** *σ* (**fixed**)), ii) step increase from spontaneous to evoked state, except for GCs (*σ* **Vary in Time**), and iii) large fixed *σ* (**Larger** *σ* (**fixed**)). The smaller and larger fixed *σ* values match the spontaneous and evoked values when *σ* varies in time. **C**) The black dots shows the region in parameter space that is within tolerance of the PSTH from data, with a 3D view (upper right) and all 2D projections. Repeated for the 3 input noise regimes. Out of 10,000 parameter sets, there are 361, 295, and 129 (respectively for **Ci, Cii, Ciii**) that are within tolerance. **D**) Blue curves show trial-averaged MC firing rate within tolerance of data (black). The gray dashed curves show the model extremes that have the worst matches to data.

**Table 1:**
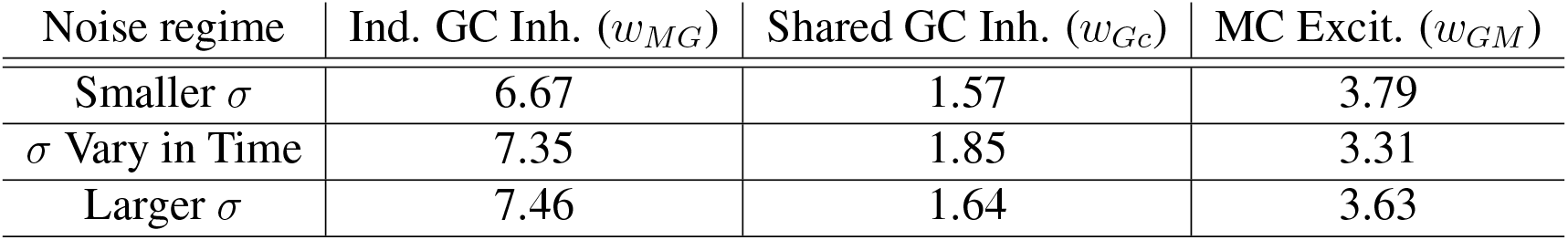
The average coupling strengths among the 10 best parameter sets in Figure 3D, model framework applied to anesthetized data yields: *w*_*MG*_ > *w*_*GM*_ > *w*_*Gc*_.

| Noise regime | Ind. GC Inh. ($w_{MG}$) | Shared GC Inh. ($w_{Gc}$) | MC Excit. ($w_{GM}$) |
| --- | --- | --- | --- |
| Smaller $\sigma$ | 6.67 | 1.57 | 3.79 |
| $\sigma$ Vary in Time | 7.35 | 1.85 | 3.31 |
| Larger $\sigma$ | 7.46 | 1.64 | 3.63 |

**Table 2:** Determining statistical significance of evoked increases in spike count variance and covariance (population average after trial-averaging), reporting the p values for two-way t-tests assuming unequal variance (Fig 5A–C). Ethyl Butyrate (**EB**), Hexanol (**HX**).

| Experimental Prep | Firing Rate | Spike Count Variance | Spike Count Covariance |
| --- | --- | --- | --- |
| Anesthetized, EB | $4.2 \times 10^{-17}$ | $5.9 \times 10^{-14}$ | $1.3 \times 10^{-66}$ |
| Anesthetized, HX | $4.1 \times 10^{-31}$ | $8.2 \times 10^{-27}$ | $4.2 \times 5.9^{-33}$ |
| Awake, EB | 0.0376 | 0.0181 | $2 \times 10^{-15}$ |
| Awake, HX | 0.154 | 0.0447 | $1.5 \times 10^{-8}$ |

**Table 3:**
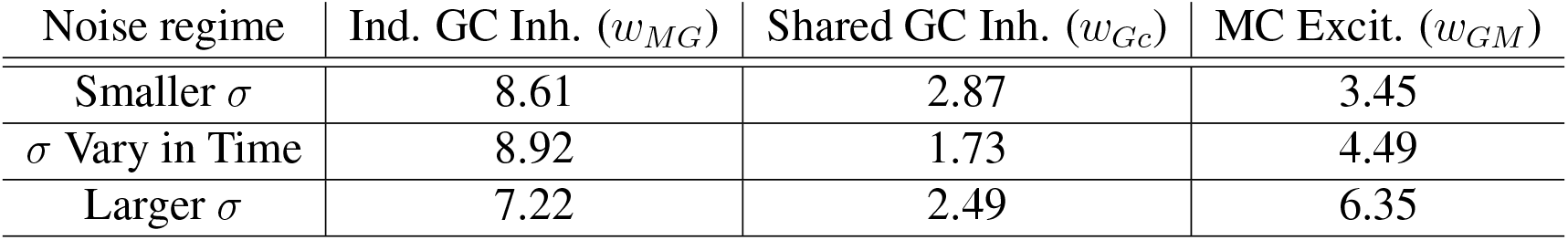
The average coupling strengths among the 10 best parameter sets in Figure 6E, model framework applied to awake data. The values are similar, having the same relationship as with anesthetized data: *w*_*MG*_ > *w*_*GM*_ > *w*_*Gc*_ (see Table 1)).

| Noise regime | Ind. GC Inh. ($w_{MG}$) | Shared GC Inh. ($w_{Gc}$) | MC Excit. ($w_{GM}$) |
| --- | --- | --- | --- |
| Smaller $\sigma$ | 8.61 | 2.87 | 3.45 |
| $\sigma$ Vary in Time | 8.92 | 1.73 | 4.49 |
| Larger $\sigma$ | 7.22 | 2.49 | 6.35 |

**Figure 3:**
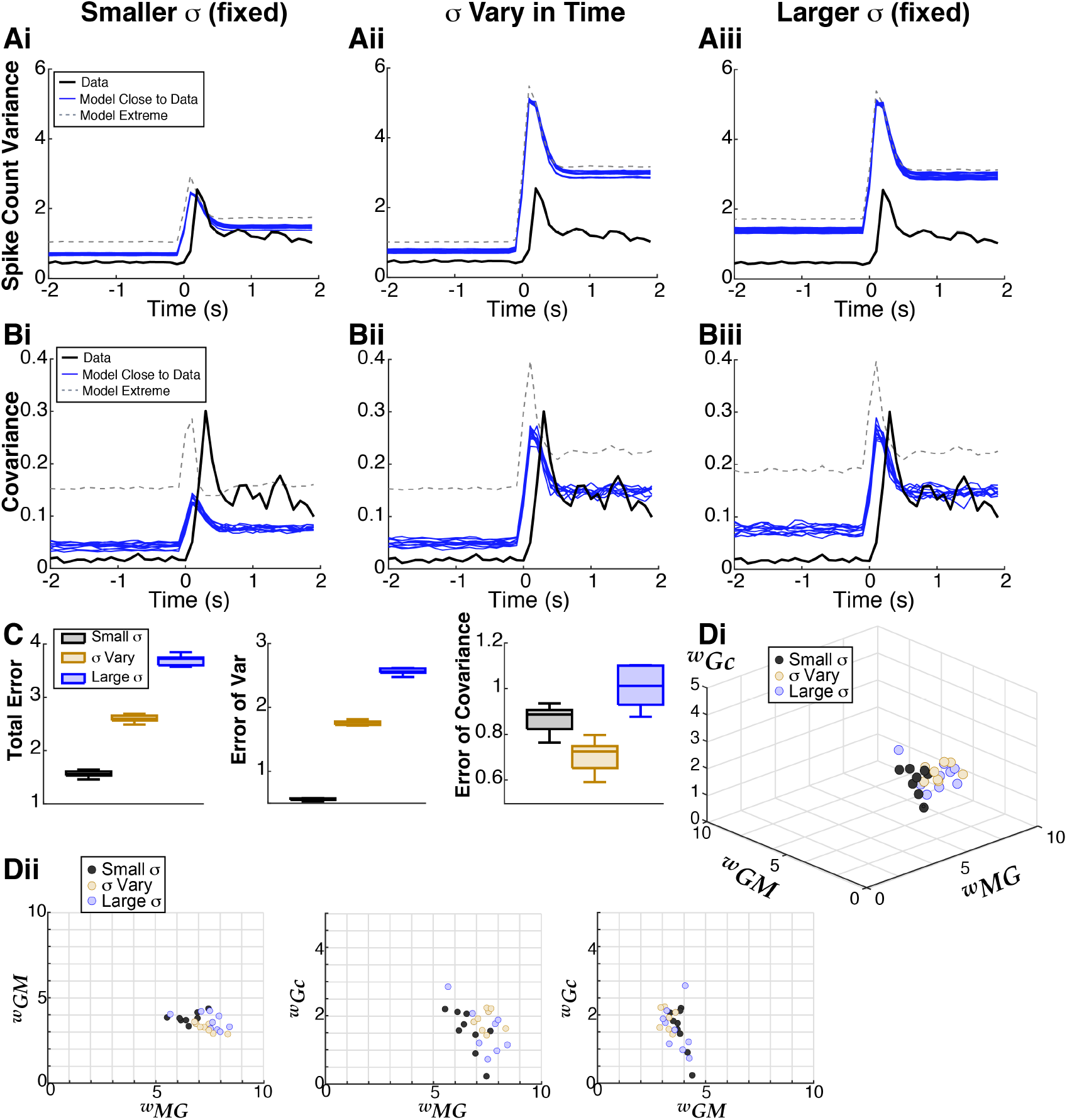
Capturing second order spike statistics with the model. Models that best match all three time-varying statistics (PSTH, spike count variance, and covariance) result in different levels of error depending on input statistics. **A**) Model results (blue) of spike count variance that best match data (black) from among set of coupling strengths that well-match the PSTH (Fig. 2C, **D**) for all 3 input noise regimes. The model extremes (gray dash) show the worst matches to the spike count variance among models that well-match the PSTH. **B**) Results for spike count covariance, same format as **A. C**) Summary of errors between model and data: small fixed *σ* (gray) result in overall best performance, followed by time-varying *σ* (tan) and large fixed *σ* (blue). Box plots for 10 best parameter sets: the shaded rectangle captures two middle quartile (25%–75%), median is solid line through shaded rectangle, and whiskers contain the most extreme points. Total Error and Error of Variance have the same ordering of input noise regimes; for matching the covariance alone, the time-varying *σ* performs best. **D**) Small region of parameter space corresponding to blue curves in **A–B. Di** shows the 3D region, **Dii** shows the 2D projections of the same points. There is some variation within a noise regime but the average values among the 10 best are very similar: see Table 1.

**Figure 4:**
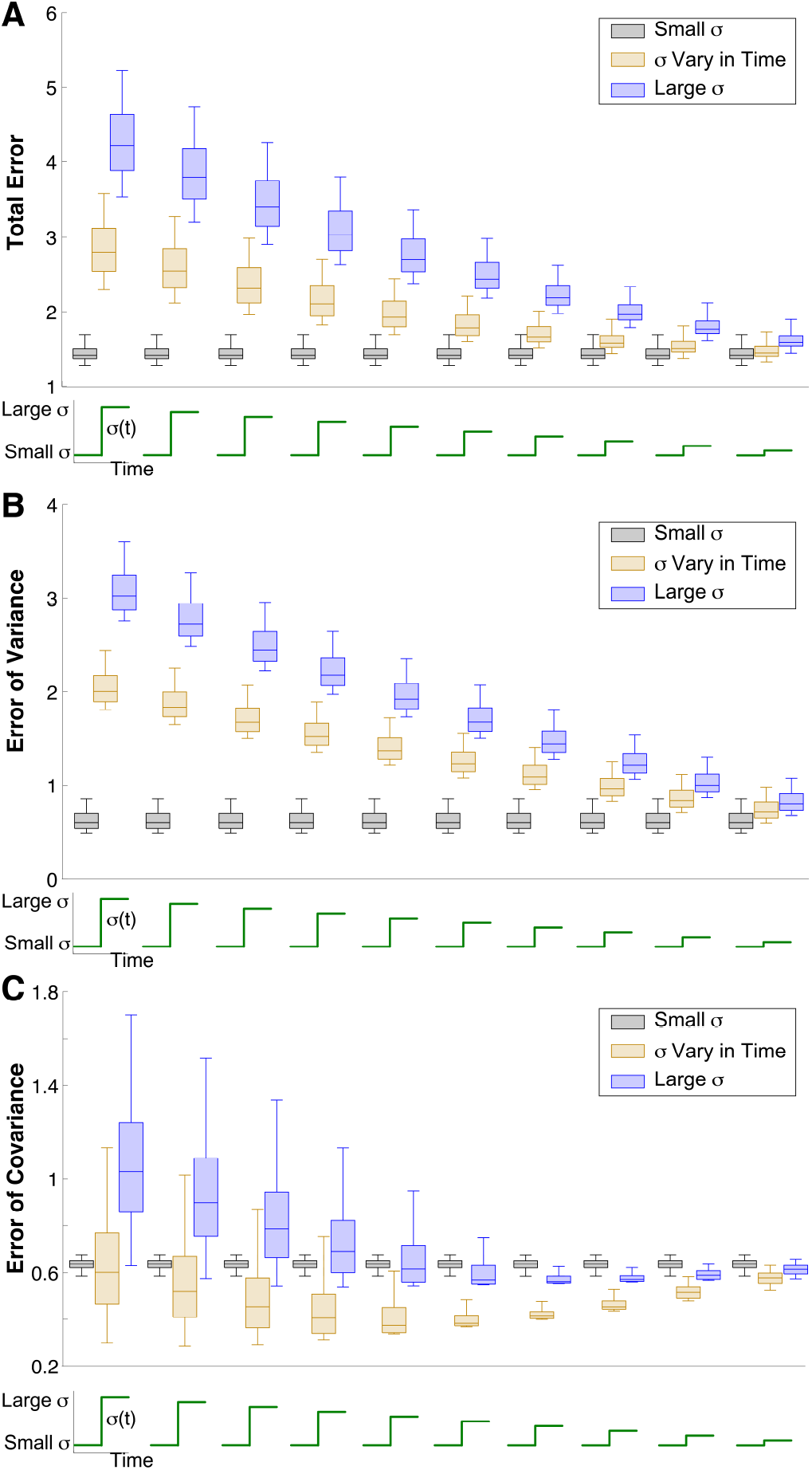
Probing other ORN input noise values. Testing other ORN input noise values to show robustness of prior results for matching time-varying data statistics, but using a steady-state approximation for computational feasibility. Each triplet of input noise values has a small fixed *σ* (black), time-varying with a step (tan), and large fixed *σ* (blue); as before, the time-varying values are set to the small values in the spontaneous state and large values in the evoked state. Results for Total Error (**A**), Error of Variance (**B**), and Error of Covariance (**C**) are all consistent with what we observed before (Fig. 3C), with the differences converging as the noise values approach each other. We tested 10 total triplets of input noise sets, using a random sample of coupling strengths (Fig. S1C) that cover the best parameter values (Fig. 3**D**), and a random set of scaling factors (Fig. S1C) to approximate spike statistics in 100 ms windows. The whiskers and inter-quartiles in the box plots capture the variation of errors over the random samples of coupling strengths and scaling factors.

**Figure 5:**
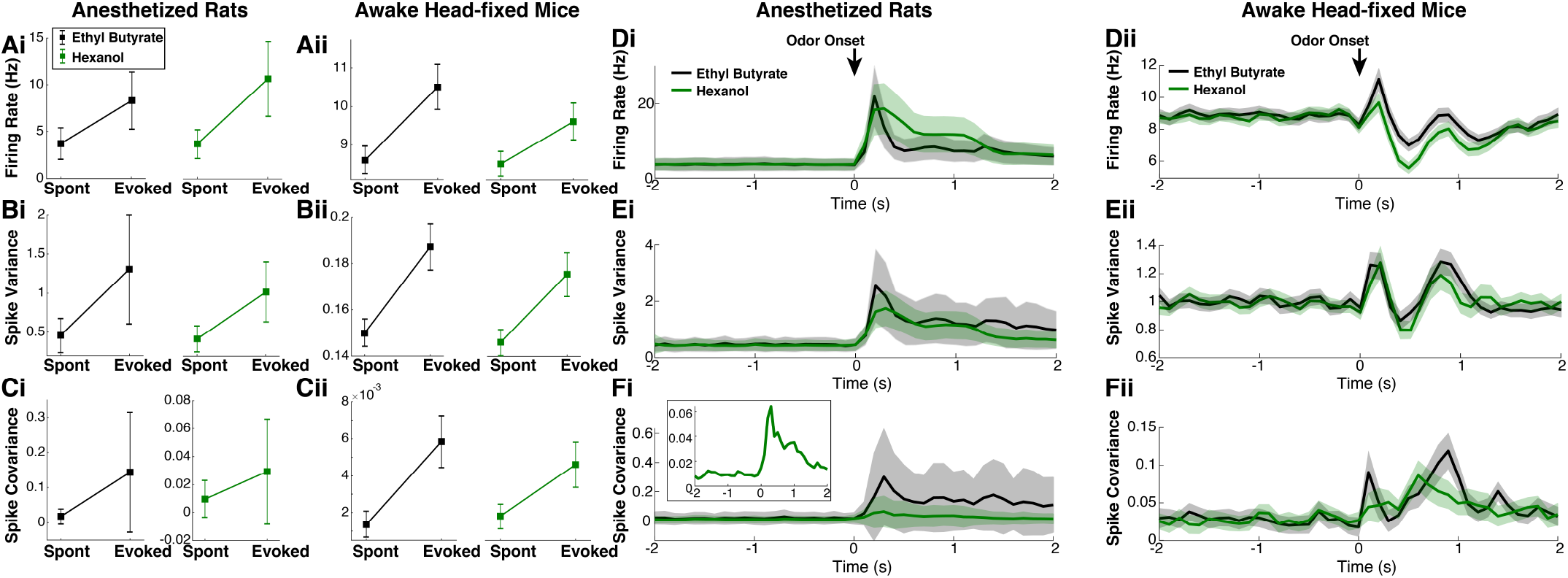
Awake breathing rodents and another odor. Anesthetized recordings (**Ai–Ci**), spike statistics generally increase in evoked state (time-averaged), with Ethyl Butyrate (**EB**, black, same as in Fig 1D–F, 1120 cells and 17674 pairs) and Hexanol (**HX**, green) with 1033 cells and 14873 pairs. Similar format to Figure 1 with squares representing population averages. With EB, percentage of cells/pairs with evoked increases: 82%, 81.1%, 57.75% (resp. for firing rate, Var, Cov); with HX, percentage of cells/pairs with evoked increases are respectively: 87.7%, 85.3%, 49.2%. Similar plots for awake (time-averaged) recordings (**Aii–Cii**) from) with same odors, 225 cells and 3203 simultaneously recorded cell pairs (15 trials per odor), with 0.3% v/v concentration, an intermediate value in Bolding and Franks (2018a). Percentage of cells/pairs increase with EB: 51.1%, 50.7%, 52.3% (resp. for firing rate, Var, Cov), percentage of cells/pairs increase with HX: 48.4%, 50.2%, 52.3%. Spontaneous and evoked periods are 200 ms before and after inhalation, much briefer because odor concentrations dissipate rapidly (*cf.* 1 s odor delivery in anesthetized) and with 20 ms time windows for more samples. Evoked increases are not as prominent on average, with less percentage of cells/pairs showing the effect. Time varying statistics in anesthetized (**Di–Fi**) with EB (same as in Fig 1), HX (green), and awake recordings (**Dii–Fii**), with firing rate (**D**), spike count variance (**E**), spike count covariance (**F**). In **A–F**, whiskers and shaded error regions are 0.2 standard deviations across population/pairs after trial-averaging for anesthetized, and 0.05 standard deviations for awake.

**Figure 6:**
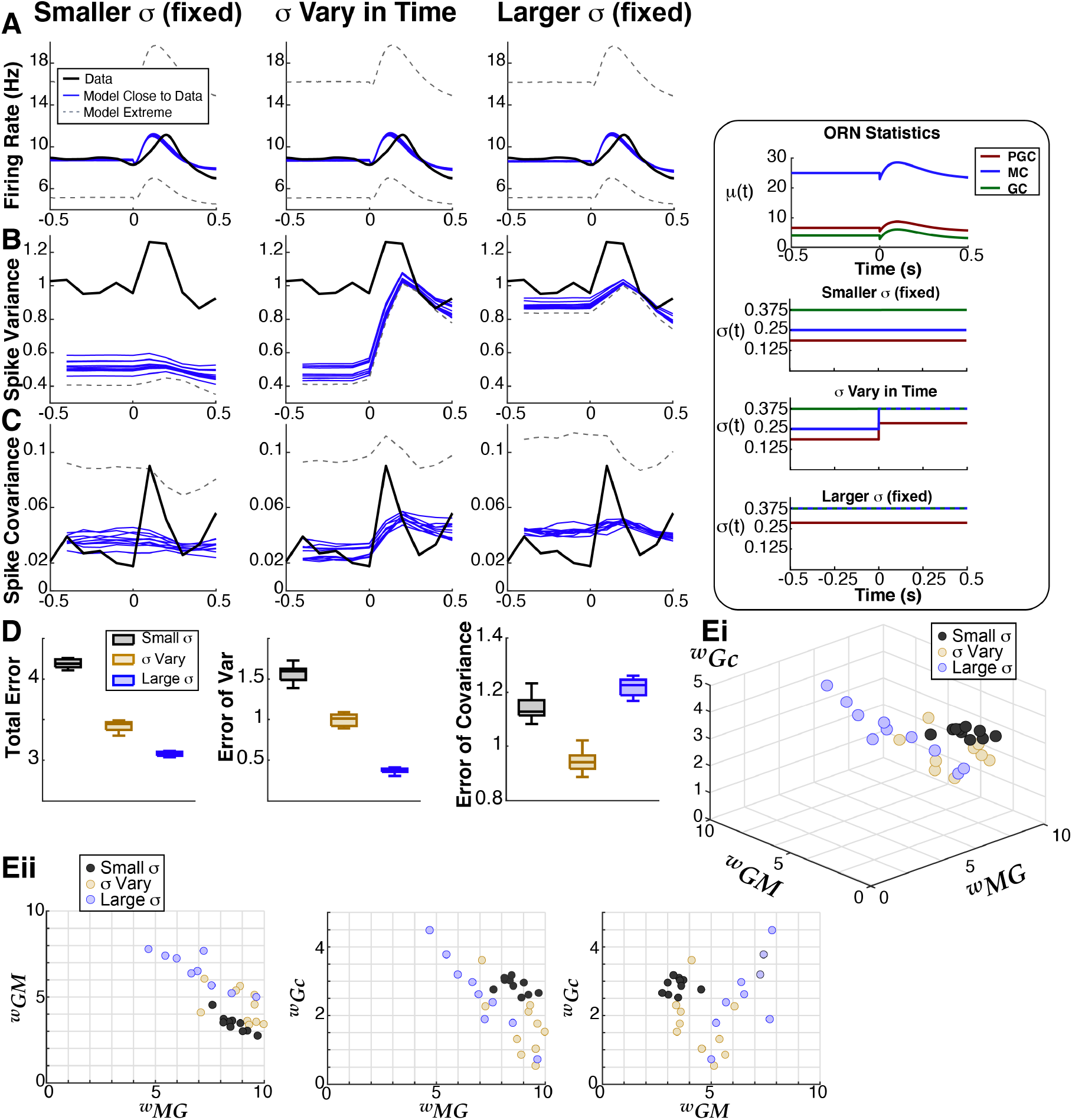
Computational model applied to awake data. Our model framework applied to awake data with EB (black curves in Fig. 5**Dii–Fii** from (2018a,b)), with analogous plots to Figures 2–3 (anesthetized data). The inset shows ORN input noise (see main text and Fig 2**B**). The model results are very similar to before when using anesthetized data: **A**) showing the best 100 fits (blue) and worse (gray dashed) to the firing rate data (black) out of the same 10,000 point parameter space. **B**)–**C**) The 10 best fits (blue) accounting for the spike variance and covariance, and the worse (gray dashed). **D**) The ORN input noise that is fixed (large here) results in overall better fits to the data, with time-varying ORN noise matching the spike covariance best. **E**) Like before, the specific coupling strengths in GC–MC subnetworks have the same relationship and similar values: *w*_*MG*_ > *w*_*GM*_ > *w*_*Gc*_ (see Table 3 for average values).

### Firing rate model

Dissecting the detailed effects of neural attributes, such as ORN inputs and OB coupling strengths, on MC spike statistic modulation is difficult experimentally, especially *in vivo*. We are currently unaware of any experiments that that simultaneously consider ORN activity and its affect on MC spiking (Scott et al. (2007) considered how ORN activity affects activity in superficial OB, but not the (significantly deeper) MC layer). Therefore we developed a simple computational model that allowed us to observe the effects of different circuit manipulations.

While realistic biophysical models can capture a lot of known electrophysiology of the mammalian OB circuit (see multi-compartment OB models by Cleland and colleagues (Li and Cleland, 2013, 2017)), they are computationally expensive to simulate. This made such models unsuitable for our study for two reasons. First, we wished to capture the time-varying statistics over a lengthy time interval of 4 seconds with 1 ms resolution, which requires many realizations in order to resolve the first and second-order firing statistics. Second, we planned to perform a robust search in a large region of parameter space. Therefore we developed a firing rate model that retains key components of the Cleland models (Li and Cleland, 2013, 2017), including the membrane time-constants and transfer functions of three important cell types (MC, GC, PGC), and the synaptic dynamics (GABA_A_, NMDA, AMPA and their associated equations, see **STAR Methods**).

With a firing rate model for each type of cell, we coupled these into a minimal sub-network of 7 cells with 2 representative glomeruli (Fig. 2**A**) each with a PGC and MC cell. The MCs receive GABA_A_ inhibition from PGC and GCs, and both PGCs and GCs receive AMPA and NMDA excitation. We also include small noise correlations that model common input from ORNs to MC (Kazama and Wilson, 2009), as well as small correlated inputs to other OB cell types (see **STAR Methods** for details). There are too many parameters in the circuit to feasibly consider, so we set the coupling strengths to be symmetric between the 2 glomeruli resulting in each MC being statistically identical, and we focus on lateral inhibition from GC and recurrent MC excitation to GC.

Our focus on lateral inhibition was informed by OB anatomy, and experimental and theoretical studies demonstrating the importance of the MC–GC network in controlling spiking co-variability. The long lateral GC dendrites are known to span multiple glomeruli and impinge on correspond MCs, thus it is reasonable to assume GC strongly impacts MC spike covariance. Giridhar et al. (2011) showed in slice recordings of OB that lateral (GC) inhibition can alter spike co-variability differently depending on if it is common or independent to MC. Lateral GC inhibition is also known to modulate MC synchrony (Galán et al., 2006). Moreover, many theoretical studies have shown the importance of correlated inhibition for modulating spiking co-variability in general circuits (Renart et al., 2010; Tetzlaff et al., 2012; Middleton et al., 2012; Ly et al., 2012). To address the role of common and independent GC inhibition, we include 3 GCs with 2 providing independent inhibition to each MC and 1 GC providing common inhibition to both MCs.

### A parameter space search reveals viable region of connection strengths

To test whether odor-evoked increases in variability stems from changes in the variability of inputs (*σ*(t)) from ORN, we considered three ways to modulate *σ*(t): i) relatively small and fixed *σ* (**Smaller** *σ*), ii) *σ* that increases in the evoked state, except for GC cells (*σ* **Vary in Time**), iii) large fixed *σ* (**Larger** *σ*), see Figure 2**B**, bottom. In the fixed input noise regimes, the smaller and larger *σ* values match the spontaneous and evoked values of time-varying *σ*(t) respectively. For all parameters, the same time-varying *μ(t)* is used (Fig. 2**B** top, see **Supplementary Material: Input statistics give reasonable model results**).

For each noise regime, we performed a robust search through 3D parameter space to find the coupling strengths (*w*_*MG*_, *w*_*GM*_, *w*_*Gc*_, see Fig. 2**A**) that match the time-varying firing rate (peri-stimulus time histogram, or **PSTH**) within a certain tolerance. Specifically, a parameter set is considered ‘Close to Data’ if: 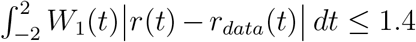, where *W*_1_(*t*) weights the uncertainty of the data (Fig. S1**A**). This tolerance results in good model fits to the data PSTH (Fig. 2**D**), and also yields hundreds of parameters for each of the 3 input noise regimes (see Fig. 2**C**).

The criteria of matching the data PSTH alone (see **STAR Methods**) drastically reduces the size of parameter space: out of 10,000 randomly distributed points, only 361 are close to the data for small and fixed input noise (Fig. 2**Ci**), 295 for time-varying input noise (Fig. 2**Cii**), and 129 for large fixed input noise (Fig. 2**Ciii**). Interestingly, the region of parameter space that results in good fits to the data PSTH looks similar for all 3 input noise regimes. The strength of synaptic input of independent GC inhibition to MC (*w*_*MG*_) is relatively large, but tends to decrease as the common GC inhibition (*w*_*Gc*_) increases in strength (lower left panels of Fig. 2**Ci, Cii**, and **Ciii**).

### Capturing MC spike variance and covariance

When we consider all 3 statistics from our data, the region of parameter space (Fig. 2C) of viable models for the PSTH should significantly shrink to reveal a more specific structure (Barreiro et al., 2017). To determine parameters that match the spike count variance and covariance, we only considered coupling strengths that are close to the data PSTH (black dots in Fig. 2C) because our goal is for a model to simultaneously match all spike statistics.

We assess how well the models match our data by:

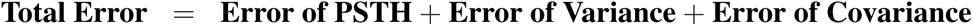

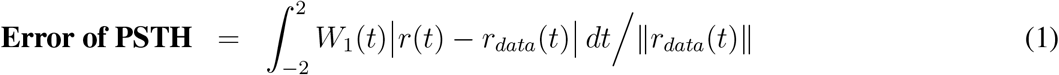

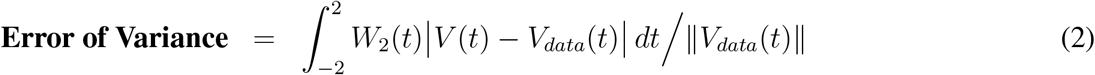

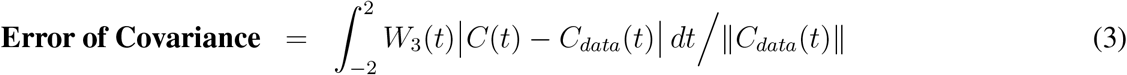

where the weights *W*_1,2,3_(*t*) account for the (population) uncertainty of the experimental data (see Fig. S1**A**), and we have normalized each integral by the magnitude of the corresponding quantity from the data 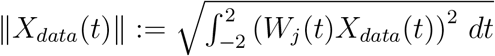.

The results are shown in Figure 3**A, B** for each of the input noise regimes. The 10 best model fits with the lowest **Total Error** are the blue curves; the parameter(s) with smallest error for both the variance and covariance are contained in the set of 10 smallest **Total Error**, for all input noise regimes. We see that the small input noise regime results in the best fit to the variance (Fig. 3**Ai**) and reasonably good fits to the covariance (Fig. 3**Bi**), but the time-varying noise fits the covariance best (Fig. 3**Bii**). The large fixed input noise regime performs the worst (Fig. 3**Aiii, Biii**). Even within the parameter sets that capture the data PSTH well, the variance and covariance can be quite different from the data (gray-dashed curves correspond to the largest error).

The errors between the model with the 10 best parameters and experimental data are shown in boxplots in Figure 3C for each of the input noise regimes. For both **Total Error** and **Error of Variance**, the small fixed input noise performs best (black), followed by time-varying input noise (tan), and large fixed input noise (blue). But for **Error of Covariance**, the time-varying input noise performs best, followed by small fixed input noise.

From Figure 3**D**, we see in all ORN input noise regimes that independent GC inhibition to distinct MC has to be large, while MC excitation is modest, and common inhibition to both MC has to be relatively weak. Although there is some variation in coupling strengths across the different noise regimes, the averages are remarkably similar (Table 1).

### Small fixed ORN input noise performs best

With a specific set of three ORN input noise values, we thus far have performed a robust search through the parameter space of coupling strengths to determine the profile of input noise that minimizes error with various statistics of our data. Next we addressed how robust these observations are by considering a range of input noise values; however, reproducing everything in the prior sections would require impractical amounts of computation. To circumvent this, we use a previously developed mathematical reduction (Barreiro and Ly, 2017a) (also see **STAR Methods: Steady-sate approximation of statistics of firing rate model**) to efficiently approximate the steady-state spiking statistics.

We used the reduction to compute statistics for a set of networks, in which we allowed the evoked input noise variance (*σ*_*large*_) to take a range of linearly spaced values, i.e., 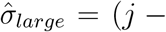 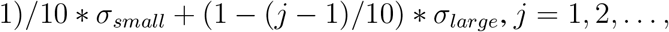 from left to right; see insets in Fig. 4). For a fair comparison of the 3 input noise regimes, we use a large random sample of the coupling strengths that maintain the same 3D structure (Fig. S1**B**), and calculated the spike statistics to get a large set of error values.

Figure 4 confirms that our results are robust. Testing a total of 10 triple input noise sets, we find that small fixed noise yields the lowest Total Error, followed by time-varying input noise and large fixed noise (Fig. 4**A**). The results for Error of Variance similarly hold (Fig. 4**B**). Finally, if matching the spike count covariance is desired, the time-varying input noise gives the smallest error (Fig. 4C).

Although our firing rate network model has only 7 cells and identical marginal MC spiking statistics, it is still sophisticated enough to exhibit odor-evoked decreases in variability (see **Supplementary Material: Firing rate model can exhibit evoked decreases in variability**, Figure S3). That is, even qualitatively matching our experimental data (i.e., increases in mean and variability at stimulus onset) requires parameter tuning in this model.

### Application to other experimental data

Here we address potential limitations of our results thus far in using our anesthetized data and a single odor. Experiments on anesthetized animals could be viewed as a limitation compared to the more natural awake state, even though complicating factors such as sniff rates and breath cycles (Wachowiak, 2011) are less significant under anesthesia. Differences in olfactory bulb activity between anesthetized and awake breathing rodents have long been known (Lowry and Kay, 2007; Kay et al., 2009), with a seminal paper by Rinberg et al. (2006) that quantified differences in MC responsiveness. Rinberg et al. (2006) found much higher spontaneous MC firing rates in awake versus anesthetized, which in principle could: i) eliminate or diminish odor-evoked increases in spiking variability, ii) vastly undermine the mechanisms our model has identified. We have thus far only considered one odor (Ethyl Butyrate, a food-like odor), whether our data observations hold with other odors has not been addressed. The (population-averaged) spike statistic comparisons with another odor, Hexanol (green curves in Figure 5) indeed shows evoked increases in spiking variability and co-variabilty. Analogous data in awake (head-fixed) mice from Bolding and Franks (2018a,b) with the same odors are shown in Figure 5Aii–Fii. Hexanol elicits different glomeruli responses than Ethyl Butyrate, but the population averaged responses are quite similar in both awake and anesthetized. The biggest difference is spike covariance in anesthetized rats, yet there is still odor-evoked increases (Fig. 5Fi, inset).

In comparison with the anesthetized data (Fig 5Ai–Ci), the odor-evoked increases are evident in all spike statistics we consider in the awake data (Fig. 5Aii–Cii), but the affects are stronger with anesthesia. The statistical significance of population average spiking increases are not as strong in awake data compared with anesthetized, see Table 2 for all p-values. The spontaneous firing rates are higher with awake mice, and the evoked firing rate changes are much smaller than in anesthetized rats (Fig. 5A,D), consistent with Rinberg et al. (2006). In fact, with Hexanol in the awake data, the evoked changes in population firing rate are minor, with a p-value of 0.154 (Table 2).

Finally, we apply our model framework to the awake data with EB and find our prior results (Figs 2–3) still hold. Because the evoked period is much shorter in awake rodents, we only consider −0.5 ≤t ≤ 0.5 s, with t = 0 the start of inhalation. The time-varying statistics return to spontaneous levels much faster than in the anesthetized data (Fig. 5Dii–Fii), and Cury and Uchida (2010) has shown that the bulk of odor information is within hundreds of ms after inhalation. In the model, there are only slight differences in the mean and variance of ORN input (see inset of Fig 6). Compared with the anesthetized setting, the mean ORN inputs here are elevated in the spontaneous state with relatively less increases in the evoked state, while the ORN variance is half as large – the resulting range of model responses encompasses the awake data.

Consistent with the model results on anesthetized data, we find the ORN input noise does not need to increase to capture the data well. The 100 best model fits, out of 10000, to the firing rate data (Fig 6) are very good for all 3 ORN noise regimes. Accounting for the spike variance and covariance (Fig 6B,C), the 10 best model fits to the data have varying levels of accuracy. Fixed ORN noise that is relatively large results in overall less total error (blue in Fig 6D and less error with spike variance, while time-varying ORN noise (tan) matches the spike covariance data best – this is what we saw in the anesthetized data (recall Fig 3C). One difference here is that when the ORN noise is small (too small), the match with the data can be worse than time-varying ORN noise. This difference is inconsequential; the conclusion that we draw from this is that ORN input noise does not need to increase to observed evoked increases in MC spike variability.

The 10 best coupling strengths in the GC–MC subnetwork that best matches the awake data have a similar relationship to what we saw before in the anesthetized data: we find large independent GC inhibition, moderate MC excitation to GC, and weak common GC inhibition: *w*_*MG*_ > *w*_*GM*_ > *w*_*Gc*_ (Fig 6E, also see Table3 for average values).

Thus, we indeed find that evoked increases in spiking variability is relatively robust, apparent in both awake and anesthetized data, but the effects are weaker in the awake mice. Applying our computational model to both awake and anesthetized data in rodents yields qualitatively similar results.

### Model prediction confirmed with awake mice

A natural prediction from our modeling results is that the ORN input noise does not have to increase in the evoked state to observe odor-evoked increases in MC spiking variability. That is, the OB circuit itself can produce increases in variability without inheriting increased variability from ORN. We tested this model prediction with a publicly available dataset (Bolding and Franks, 2018b) originally collected for Bolding and Franks (2018a). The Franks lab performed direct optical MC stimulation with 1 s light pulses in awake and freely breathing mice (Fig. 7 in (Bolding and Franks, 2018a)). By circumventing the normal sensory pathway with direct MC stimulation, the ORN input statistics (i.e., variance) presumably remains similar during light-evoked MC spiking. Hence our model prediction would be confirmed if the spike count variance and covariance increases with optogenetic stimulation.

**Figure 7:**
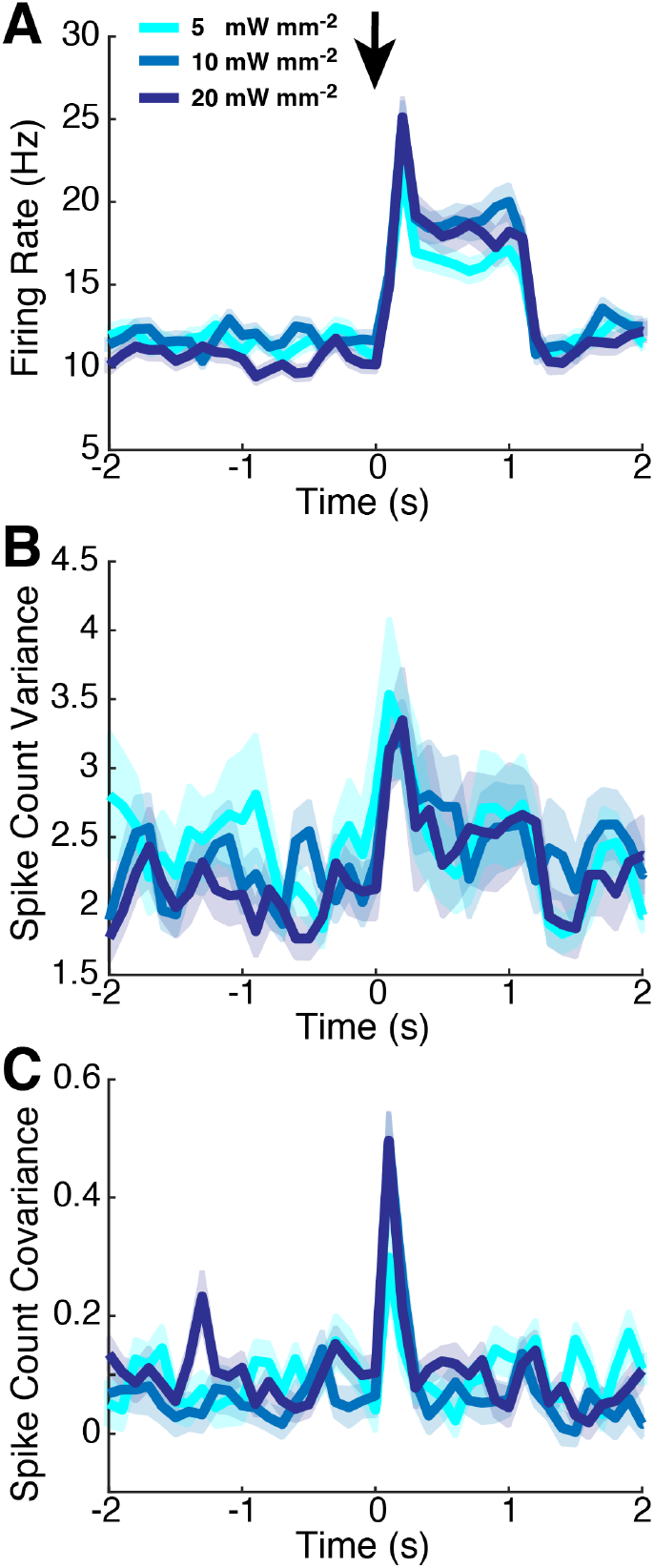
Data from the Franks lab shows evoked increases in MC spiking variability. Using publicly available data (Bolding and Franks, 2018b), we verify our model prediction that ORN input noise does not have to increase to observe evoked increases in MC spiking variance and covariance. We use the same 3 light intensities as in Bolding and Franks (2018a). Population-averaged statistics displayed here are **A**) PSTH, **B**) spike count variance, and **C**) spike count covariance. The PSTH increases in the evoked state, and the spike count covariance clearly increases for a brief period after light stimulation. Evoked increases of spike count variance is harder to see, but two-sample t-test assuming unequal variance shows statistically significant increases for all light intensities (*p* – value = 0.0296, 5.5 × 10^−4^, 1.2 × 10^−4^, respectively). Spike counts were calculated in 100 ms half-overlapping windows; there were 77 total cells and 812 total pairs, with 10 trials for each light intensity. Shaded regions show relative population heterogeneity after trial-averaging: μ ± 0.05std (standard deviation across population/pairs after trial-averaging).

Figure 7 indeed shows evoked increases in OB spike count variance (Fig. 7**B**) and covariance for several light intensities (same ones used in Bolding and Franks (2018a)). The OB spike covariance clearly increases after light stimulation (Fig. 7C), although the increases is brief, persisting for only about 300 ms. The spike variance, likewise, increases after light stimulation (Fig. 7**B**). This increase is less noticeable by visual inspection (than the increases visible in 7**A,C**), but two-sample t-tests assuming unequal variance confirm that the increase from spontaneous to evoked is statistically significant with *p*–values of 0.0296, 5.5 × 10^−4^, 1.2 × 10^−4^ for the three respective light intensities (5, 10, 20 mW/mm^2^).

## Discussion

Using recordings of spike activity in OB of anesthetized rat and awake mice we demonstrate that variability and co-variability of spiking increases at stimulus onset. This phenomenon is relatively understudied, in contrast to the stimulus-induced decreases in (co-)variability that have been widely reported in many cortical and motor regions (Churchland et al., 2010; Doiron et al., 2016). Using a biophysically-inspired model, we studied the mechanisms of stimulus-induced increases in spiking variability and co-variability in OB. We have described in detail how both ORN input variability and interactions within the OB circuit may play a role. We verify the importance of OB circuit interactions using previously reported data from optogenetic stimulation in OB of awake mice (Bolding and Franks, 2018a,b), which provides OB stimulation while holding ORN input variability fixed.

Our findings complement previous reports of odor-evoked increases in (co-)variability of OB cells in insects and fish (Kazama and Wilson, 2009;Wanner and Friedrich, 2020), establishing this phenomenon in mammals. Recent recordings in the OB of developing zebrafish show time-varying statistics of correlation that are similar to what we observed in rat: the correlation of activity measured by calcium imaging increased suddenly in the evoked state, followed by a slow decay to a level that remained above the spontaneous values (Wanner and Friedrich, 2020). These authors explained correlation of mitral cells by analyzing detailed connectivity structures of interneurons and MCs using electron microscopy, neglecting GCs because they have an insignificant role at this stage of zebrafish development. Kazama and Wilson (2009) also showed that the correlation of projection neurons (**PN**, analogous to MC) in *Drosophila* OB increased in the evoked state compared to spontaneous. This was observed for pairs of PNs from the same or different glomeruli, and independent of whether ORN inputs were shared or not. They attributed the increase in PN spike correlation primarily to common inputs from ORN rather than lateral connections in the glomeruli layer. We focussed on both ORN inputs and MC–GC coupling strengths because they likely have significant affects on MC spike dynamics in mammals (Cang and Isaacson, 2003; Galán et al., 2006; Giridhar et al., 2011). Taken together, our results and these previous studies suggest that evoked increases in spiking variability in the OB seems to be a core principle, generalizing across many species.

One important prediction we make here is that evoked increases in MC spike count variability can occur without increases in the noise of ORN input. We verified this prediction with another dataset (Bolding and Franks, 2018a,b), in which direct optical stimulation was used to bypass the normal sensory pathway, thus avoiding an increase in sensory input variability. To our knowledge, whether ORN spiking variance actual stays constant in natural conditions has not been directly verified. There have been many ORN recordings in mammals (Spors et al., 2006), *Drosophila* (Nagel and Wilson, 2011; Kazama and Wilson, 2009) and locusts (Raman et al., 2010), but the trial-to-trial variability of ORN spiking in both spontaneous and evoked states has not been reported or is not readily available. Duchamp-Viret et al. (2005) characterized the noisy spiking of ORN in tracheotomized and breathing rats but only in the spontaneous state. The data in Figure 1 of Nagel and Wilson (2011) has shaded gray regions to show trial-variability of ORN spiking in *Drosophila*, and indeed by visual inspection the trial-variability appears to be constant in spontaneous and evoked states for several different odors (but not all), consistent with our model prediction. Although in our model the net effect of ORN inputs and computation at the glomeruli level is combined, glomeruli processing separately could also impact increases in MC spike variability.

ORN responses are known to have odor-evoked increases in spatiotemporal variability, for example in locusts (Raman et al., 2010) and mammals (Spors et al., 2006). Calcium imaging of ORN inputs to OB in anesthetized breathing rats *in vivo* has shown that across glomeruli, the calcium activity with the same odor we use (ethyl butyrate) can vary dramatically compared to the spontaneous state (Spors et al., 2006). Although our OB model consists of only 2 glomeruli and is far from a realistic spatial model, these data appear to be at odds with our model prediction that ORN input variance is similar in both states. However, Spors et al. (2006) also showed that higher odor concentrations may result in less variability across glomeruli (their Fig. 2B).

We also predict the relative strength of three important coupling strengths within OB: independent GC inhibition to distinct MCs (*w*_*MG*_), shared GC inhibition to MCs (*w*_*Gc*_), and MC excitation of GCs (*w*_*GM*_). We found that shared GC inhibition was significantly weaker than independent GC inhibition (about one-quarter of the magnitude on average). The strength of MC excitation was intermediate, stronger than shared GC inhibition but weaker than independent GC inhibition. The coupling strength values that best matched the data were very similar for across different ORN input variance types (small fixed, large fixed, and variable in time). We are unaware of any experiments that refute or validate these predictions about coupling strengths within OB, but relatively strong GC inhibition is well-known in the mammalian OB (Cang and Isaacson, 2003; Schoppa and Urban, 2003; Boyd et al., 2012; Markopoulos et al., 2012; Burton and Urban, 2015). Nevertheless, we hope our model predictions about OB synaptic strengths will be insightful in further mechanistic studies of OB spiking modulation.

### Limitations of the study

We conclude by listing several factors which could in principle affect our results, but which we were not able to consider here. Our modeling did not account for all common odors or odor mixtures, although the MC responses with both Hexanol and Ethyl Butyrate (in anesthetized rats and awake mice) have qualitatively similar population responses (Fig. 5). Our modeling results are consistent with both anesthetized and awake data, but we did not dissect the awake data into different sniff frequencies that can alter temporal neural activity Cury and Uchida (2010). The OB model did not explicitly model feedback from piriform or other cortical areas even though there is evidence of odor-specific feedback from piriform that can decorrelate OB responses (Otazu et al., 2015). Different odors elicit different spatiotemporal OB responses, and whether our modeling results would hold qualitatively with other odors beyond EB and HX is an open question, but we note that evoked increases in spike variability in our data is consistent with awake mice and under optogenetic stimulation Bolding and Franks (2018a), as well as with other species with other odors, including *Drosophila* (Kazama and Wilson, 2009), zebrafish (Wanner and Friedrich, 2020). Also, in our models we fixed the transfer functions (Fig. 2**A**) for computational tractability, but note that varying this component of the model could lead to different spike statistic dynamics (de la Rocha et al., 2007; Tetzlaff et al., 2012; Barreiro and Ly, 2017b, 2018) (also see Hiratani and Latham (2020)).

In our model, we only surveyed GC/MC subnetwork parameters in the OB circuit because GCs outnumber other cell types and they are known to span multiple glomeruli (i.e., strongly modulate MC co-variability). We kept the coupling strengths of PGC/MC subnetworks fixed, even though inhibition from PGC (Wanner and Friedrich, 2020) and other interneurons (Burton, 2017) can play a key role in modulating MC spiking. Specifically, PGC activity could lead to parvalbumin (PV) interneurons strongly inhibiting MC (and tufted cells), an effect we did not explicitly model. Our array recordings in the MC layer detected spikes from putative MCs (similarly in Bolding and Franks (2018a) that we have included), thus accounting for differences in MC and tufted cell firing would require incorporating another dataset. Others have shown that MC and tufted cell firing rates can defer and may have different roles in relaying odor information (Geramita et al., 2016). Also, recent work has shown that tufted cells and MCs fire at different phases of the breath cycle (with MC having less precision) (Fukunaga et al., 2012). PGC inputs differentially effect MC and tufted cells, causing this delay (Fukunaga et al., 2014). Considering the effects of PGC on MC spiking in detail is absent from our modeling work here. Moreover, we chose to apply the same model framework to both awake and anesthetized datasets even though a more accurate model of the awake state would account for responses modulating with the breath cycle. The model fits to the anesthetized data (Figs. 2–3) are generally better than with the awake data (Fig. 6); also, the best coupling strengths in the GC–MC are confined to a smaller region in the anesthetized model results. Whether an awake model accounting for the breath cycle could account for these different model results was unexplored by us. Despite these limitations, we have demonstrated the utility of our model by verifying a prediction in awake mice.

We use a network model that does not incorporate heterogeneity of MCs or other cells, so the model cannot simultaneously match the 3 statistics we focus on *as well as* the Fano factor and spike count correlation. Fano factor and correlation in our data are not simply scaled by the population mean and variances (as they would be in a homogeneous network model) because of the large degree of heterogeneity. All cells/pairs do not simply increase or decrease uniformly in the evoked state (Fig. 1**D–F**), and the modulation of (co-)variability could strongly depend on the underlying function or selectivity of individual cells/pairs (Ruff and Cohen, 2014; Moreno-Bote et al., 2014). Finally, limiting our investigation to a single odor means that we did not explicitly consider information or coding: much theoretical work has been devoted to whether relatively larger co-variability is better (da Silveira and Berry II, 2014) or worse (Ecker et al., 2011) for coding, with a recent study showing this also depends on population size (Bartolo et al., 2020). This intersects with the question of heterogeneity, because the relationship between collective population statistics (i.e., not just population-averaged statistics) and efficient/accurate neural coding is complicated (Kohn et al., 2016). We hope to consider each of these important issues in future work.

## STAR Methods

### RESOURCE AVAILABILITY

#### Lead contact

Further information and requests for code and data should be directed to and will be fulfilled by the lead contact Cheng Ly

#### Materials availability

This study did not generate new unique reagents.

#### Data and code availability

See https://github.com/chengly70/OB for MATLAB code implementing all models in this paper, as well as code to display the Franks’ lab data and our data.

See https://www.dx.doi.org/10.6084/m9.figshare.14877780 for anesthetized data generated by the Shew Lab.

### EXPERIMENTAL MODEL AND SUBJECT DETAILS

#### Electrophysiological recordings

All procedures were carried out in accordance with the recommendations in the Guide for the Care and Use of Laboratory Animals of the National Institutes of Health and approved by University of Arkansas Institutional Animal Care and Use Committee (protocol #14049). Data were collected from 11 adult male rats (240-427 g; *Rattus Norvegicus*, Sprague-Dawley outbred, Harlan Laboratories, TX, USA) housed in an environment of controlled humidity (60%) and temperature (23°C) with 12 h light-dark cycles. The experiments were performed in the light phase.

##### Surgical preparations

Anesthesia was induced with isoflurane inhalation and maintained with urethane (1.5 g/kg body weight (**bw**) dissolved in saline, intraperitoneal injection (**ip**)). Dexamethasone (2 mg/kg bw, ip) and atropine sulphate (0.4 mg/kg bw, ip) were administered before performing surgical procedures. Throughout surgery and electrophysiological recordings, core body temperature was maintained at 37°C with a thermostatically controlled heating pad. To isolate the effects of olfactory stimulation from breath-related effects, we performed a double tracheotomy surgery as described previously (Gautam and Verhagen, 2012). A Teflon tube (OD 2.1 mm, upper tracheotomy tube) was inserted 10 mm into the nasopharynx through the rostral end of the tracheal cut. Another Teflon tube (OD 2.3 mm, lower tracheotomy tube) was inserted into the caudal end of the tracheal cut to allow breathing, with the breath bypassing the nasal cavity. Both tubes were fixed and sealed to the tissues using surgical thread. Local anesthetic (2% Lidocaine) was applied at all pressure points and incisions. Subsequently, a craniotomy was performed on the dorsal surface of the skull over the right olfactory bulb (2 mm x 2 mm, centered 8.5 mm rostral to bregma and 1.5 mm lateral from midline).

##### Olfactory Stimulation

A Teflon tube was inserted into the right nostril and the left nostril was sealed by suturing. Odorized air was delivered for 1 s in duration at 1 minute intervals. The odorant was Ethyl Butyrate (**EB**) and Hexanol (**HX**), both at saturated vapor. We note that the full experimental data set has retronasal stimulation, but here we consider only orthonasal stimulation.

##### Electrophysiology

A 32-channel microelectrode array (**MEA**, A4×2tet, NeuroNexus, MI, USA) was inserted 400 *μ*m deep from dorsal surface of OB targeting tufted and mitral cell populations. The MEA probe consisted of 4 shanks (diameter: 15 *μ*m, inter-shank spacing: 200 *μ*m), each with eight iridium recording sites arranged in two tetrode groups near the shank tip (inter-tetrode spacing: 150 *μ*m, within tetrode spacing 25 *μ*m). Simultaneous with the OB recordings, we recorded from a second MEA placed in anterior piriform cortex. Voltage was measured with respect to an AgCl ground pellet placed in the saline-soaked gel foams covering the exposed brain surface around the inserted MEAs. Voltages were digitized with 30 kHz sample rate (Cereplex + Cerebus, Blackrock Microsystems, UT, USA). Recordings were band-pass filtered between 300 and 3000 Hz and semiautomatic spike sorting was performed using Klustakwik software, which is well suited to the type of electrode arrays used here (Rossant et al., 2016).

### METHOD DETAILS

#### Firing rate model details

Our firing rate model contains many features of a multi-compartment biophysical OB model (Li and Cleland, 2013, 2017), including the synaptic variables, transfer functions, and membrane time constants. But this model is also simple enough to enable fast simulation times to assess how various attributes modulate firing statistics; see Kanashiro et al. (2017) for similar use of firing rate models. The voltage, intrinsic currents and all associated dynamic variables are modeled with a single state variable *x*_*j*_ that is a phenomenological representation of the cell’s activity, which is mapped to a firing rate via a monotonically increasing function *F*_*j*_.

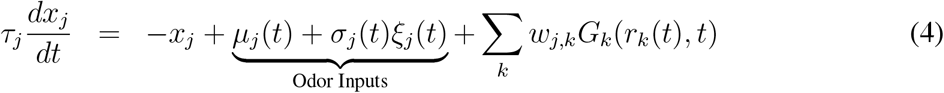

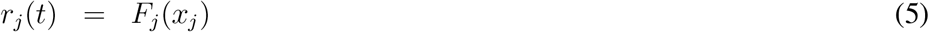

Noise terms *ξ*_*j*_(*t*) represent inputs, are white in time (i.e. ⟨*ξ*_*j*_(*t*)⟩ = 0, ⟨*ξ*_*j*_(*t*) *ξ*_*j*_(*t*′)⟩ = *δ*(*t* - *t*^′^)), and are pairwise correlated (Galán et al., 2006; Marella and Ermentrout, 2010; Litwin-Kumar et al., 2011). Input noise terms are correlated between an MC and PGC pair within a glomerulus because they have input connections from the same ORN cells (⟨*ξ*_*j*_(*t*)*ξ*_*k*_(*t*)⟩ = 0.1), correlated between the two MCs because of common ORN inputs (Kazama and Wilson, 2009) (⟨*ξ*_*j*_(*t*)*ξ*_*k*_(*t*)⟩ = 0.05), and between the GCs (⟨*ξ*_*j*_(*t*)*ξ*_*k*_(*t*)⟩ = 0.05) because they are known to synchronize (Li and Cleland, 2017; Marella and Ermentrout, 2010). The correlation values are small (0.05 or 0.1), with the largest value between PGC and MC (0.1) corresponding to common ORN inputs (Kazama and Wilson, 2009). All other pairs of cells have no input correlation.

The transfer functions *F*_*j*_(·) (Fig. 2**A**) we use is numerically calculated from our implementation of the Cleland models (Li and Cleland, 2013, 2017). For simplicity, we choose static functions *F*_*j*_ for each cell type that is similar to several computed transfer functions, see **Supplementary Material: Details of the transfer function calculation** Figure S4C. Any inputs outside of the ranges in Figure 2A are capped, so inputs below the input range give 0 firing and inputs larger than the maximum input range result in the maximum firing rate.

We use the synapse equations from the Cleland models but replace the presynaptic voltage with a function of firing rate *f*(*r*(*t*)) since firing rates are what drive average synaptic activity:

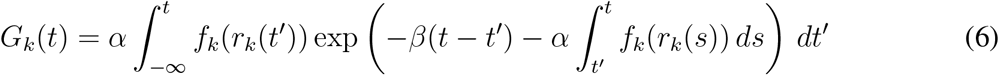

Here the strengths of the synapse *w*_*j,k*_ is a signed-quantity: < 0 for GABA_A_ and > 0 for both AMPA and NMDA; but all plots of coupling strengths (*w*_*MG*_, *w*_*Gc*_) show the absolute value. The parameters *α*, *β* are obtained directly from the biophysical model (Li and Cleland, 2013, 2017) and vary with synapse type (i.e., α_*NMDA*_). Here we choose: 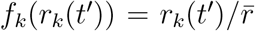 where 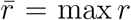. The synaptic filter 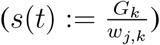 satisfies the synapse equations used in the biophysical model (20) but with *a*_*on*_(*v*_*pre*_) replaced with *f*(*r*(*t*)). The steady state of *G*_*k*_ is:

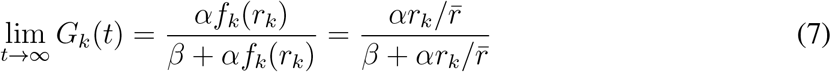

The spike count variance/covariance is modeled by the variance/covariance of the firing rate integrated over a time window *T*. That is, we define the random variable

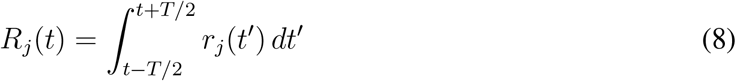

and use *Var*(*R*_*j*_) and *Cov*(*R*_*j*_, *R*_*k*_) as proxies for the spike count variance and covariance respectively. For this study, we focused on a time window of *T* = 100 ms.

Each network was simulated using the explicit Euler-Maruyama method for stochastic differential equations with a resolution of 1 ms, for a simulation time of 4 seconds (2 seconds before stimulus onset and 2 seconds afterwards). For each set of parameter values, we simulated 100,000 realizations in order to accurately resolve the mean value of second-order statistics such as spike count covariance.

#### Coupling strength parameter space and error calculations

For each ORN noise regime, we calculate model PSTH *r*(*t*) for 10,000 points that fill the 3D volume of parameter space generated by Halton sequence sampling (Kocis and Whiten, 1997). To account for uncertainty about a population-averaged statistic, we weight the difference between the model and data PSTH *r*_*data*_(*t*) with *W*_1_(*t*), which is a monotonically decreasing function of the standard deviation (across cells) of the data PSTH (Fig. S1**A**). Error calculations for the spike count variance and covariance are computed analogously.

#### Steady-state approximation of statistics of firing rate model

We derived a steady-state approximation of the complete set of 1^st^ and 2^nd^ order statistics of a general firing rate model (i.e., equations (4)–(5)), with a weak coupling assumption in another paper (Barreiro and Ly, 2017a). The approximation consists of a set of transcendental equations that are to be solved self-consistently; this procedure is more computationally efficient than performing Monte Carlo simulations and averaging over realizations. Here we augment that method to allow for larger coupling strengths that are necessary for capturing the spike statistics of our experimental data as follows. We use a new set of mean parameter values that is shifted from *μ*_*j*_, resulting in a new operating point where the network coupling is effectively weak. The shift in the *μ*_*j*_ parameters is accomplished in a principled manner and is different depending on the coupling strengths *w*_*j,k*_ and *μ*_*j*_’s. We replace *μ*_*j*_ with:

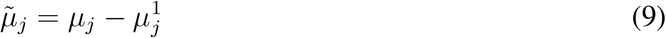

where 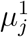 is the solution to:

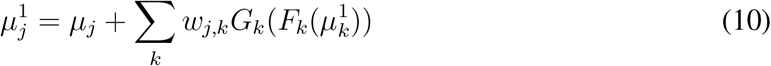

Then we find a solution to the following set of equations:

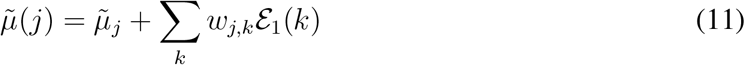

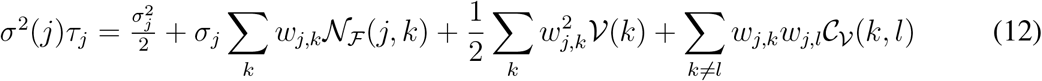

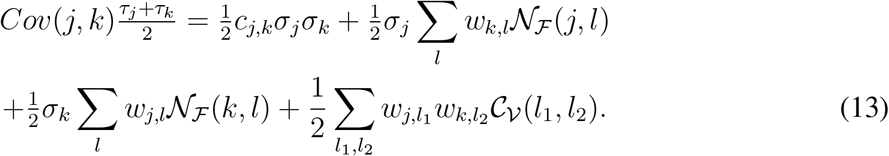

where the definitions of 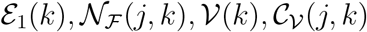 are in Table 4.

**Table 4:** Definitions of variables for the steady-state approximation of statistics (11)–(13). Whenever *j* = *k* in the double integrals (e.g., in 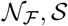), the bivariate normal distribution *ϱ*_*j,k*_ is replaced with the standard normal distribution *ϱ*_1_. Note that order of the arguments matters in 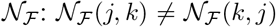 in general; all of these quantities depend on 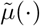, *σ*(·). For notational convenience, we define *H*_*k*_(*x*) ≔ *G*_*k*_ (*F*_*k*_(*x*)).

| Abbreviation | Definition |
| --- | --- |
| $\mathcal{E}_1(k)$ | $\int H_k(\sigma(k)y + \tilde{\mu}(k)) \varrho_1(y) dy$ |
| $\mathcal{E}_2(k)$ | $\int H_k^2(\sigma(k)y + \tilde{\mu}(k)) \varrho_1(y) dy$ |
| $\mathcal{V}(k)$ | $\int H_k^2(\sigma(k)y + \tilde{\mu}(k)) \varrho_1(y) dy - \left( \int H_k(\sigma(k)y + \tilde{\mu}(k)) \varrho_1(y) dy \right)^2$<br>$= \mathcal{E}_2(k) - [\mathcal{E}_1(k)]^2$ |
| $\mathcal{N}_{\mathcal{F}}(j, k)$ | $\iint H_k(\sigma(k)y_1 + \tilde{\mu}(k)) \frac{y_2}{\sqrt{2}} \varrho_{j,k}(y_1, y_2) dy_1 dy_2$, if $j \neq k$<br>$\int H_j(\sigma(j)y + \tilde{\mu}(j)) \frac{y}{\sqrt{2}} \varrho_1(y) dy$, if $j = k$ |
| $\mathcal{S}(j, k)$ | $\iint H_j(\sigma(j)y_1 + \tilde{\mu}(j)) H_k(\sigma(k)y_2 + \tilde{\mu}(k)) \varrho_{j,k}(y_1, y_2) dy_1 dy_2$ |
| $\mathcal{C}_{\mathcal{V}}(j, k)$ | $\mathcal{S}(j, k) - \mathcal{E}_1(j)\mathcal{E}_1(k)$ |

The final step in our approximation is to shift the mean back: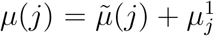 and solve for the firing statistics:

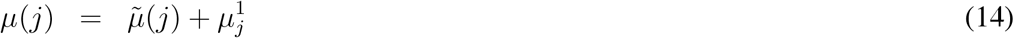

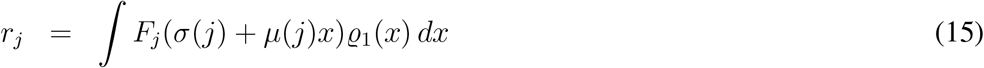

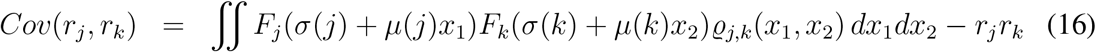

Where *ϱ*_1_ is the standard normal distribution and *ϱ*_*j,k*_ is a standard normal bivariate distribution with correlation 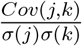 from Eqs. (12)–(13).

We can compare the firing rate *r*_*j*_ (Eq. (15)) directly to the corresponding value from our experimental data, but the approximate variance and covariance (Eq. (16)) are computed for instantaneous time rather than the 100 ms time windows. Unlike firing rate, second-order statistics can depend on the time window used for measurement in nontrivial ways (Gabbiani and Koch, 1998). To model spiking activity in 100 ms windows, we hypothesized that the relationship between statistics measured at instantaneous time and statistics measured in 100 ms bins could be modeled by a scaling factor. We used results from Monte Carlo simulations of the firing rate model (Eqs. (4)–(8)) to find this scaling factor as described below.

We computed the ratio between variances and covariances from the steady-state approximation (variances and covariances, in either the spontaneous or evoked state), and the corresponding statistic in each 100 ms bin from the 30 best parameter simulations (10 each for small *σ*, varying *σ*, and large *σ*). For example, 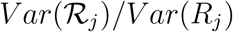 gives us a variance ratio for cell j, where R_*j*_ are spike counts in 100 ms windows (Eq. (8)) and 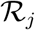 are spike counts in 1 ms windows 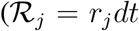, see Eq. (5)). Similarly 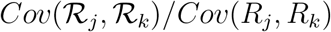 gives us a covariance ratio for cells j, k. This yielded a range of values in the vicinity of 2±0.5, with variation between simulations, across time, and across statistics (variance vs. covariance, see Fig. S1C). However, we did notice a correlation between ratios for variance and covariance involving the same cells; therefore we grouped statistics into pairs, e.g. 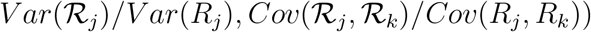. We show these paired ratios as black dots in each panel of Figure S1C; superimposed we show the time-averaged values during spontaneous and evoked activity in red, tan and blue, respectively.

These panels demonstrate several points about the ratio between instantaneous and 100 ms time bin statistics: first, for a given cell or cell pair the ratios for variance and covariance are correlated; second, the ratios vary even within a single simulation; third, all of the ratios have the same order of magnitude and lie within 25% of an average value of 2. Thus, when we measure error for the networks for which only the instantaneous rates have been computed (i.e. Figure 4), we will actually use a large number of scaling factor pairs to produce a distribution of error values. Each is plausible, given the uncertainty in our “fudge factor”. Since the ratios are fairly similar, we use the same random sample of 500 ratios (black dots) for all noise regimes; see ‘get RatiosRand.m’ at https://github.com/chengly70/OB.

Finally, we generate a random set of 1000 coupling strengths for each noise regime (black dots in Fig. S1**B**) as follows. For a given noise regime, we take the 10 best sets (red, tan or blue in Fig. S1**Bi–Biii**), calculate the mean of (*w*_*GM*_, *w*_*MG*_, *w*_*Gc*_), denoted by 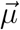 and their 3 × 3 covariance matrix Cw. We then calculate a Cholesky decomposition Cw = **LL**^′^ (**L** is a 3 × 3 lower triangular matrix), and generate 990 multivariate normal random variables:

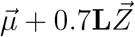

where 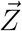 is an instance of a 3 × 1 uncorrelated standard normal. The samples have the same mean (*w*_*GM*_, *w*_*MG*_, *w*_*Gc*_) (up to finite sampling error) and similar covariance (compare black dots with red, tan and blue in Fig. S1**B**). The 990 random samples are combined with the 10 best coupling strengths for a total of 1000 sets; see ‘get GsRand.m’ at https://github.com/chengly70/OB.

So for each noise regime (i.e., each error bar in Fig. 4, there is a large set of errors between experimental data and model that stem from both the random sample of coupling strengths (1000 sets, black dots) and random ratios (500 sets, black dots). With a steady-state approximation, we must augment the error calculation; we use the same weights as before (see Fig. S1**A**) except we only consider a 1 s period in the spontaneous state (−2 ≤ *t* ≤ −1) and a 1 s period in the evoked state (1 ≤ *t* ≤ 2).

## QUANTIFICATION AND STATISTICAL ANALYSIS

Statistics of anesthetized data from the Shew Lab and Franks Lab were calculated by computing the trial average statistics for each cell/pair, then taking the population average. Spike statistics were calculated with half-overlapping time windows so that each time bin has 3 instances of trial data. Statistical significance was calculated using two-way t-tests assuming unequal variance using the MATLAB function ttest2(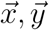,’VarType’,’unequal’). Methods are provided in: https://github.com/chengly70/OB/tree/master/ProcessShewData and https://github.com/chengly70/OB/tree/master/ProcessFranksData, respectively, using built-in MATLAB functions.

Statistics of the firing rate model were calculated analogously but without taking a population average because the network is homogeneous. Methods are provided in: https://github.com/chengly70/OB/tree/master/RateModel.

## ADDITIONAL RESOURCES

See Supplementary Material for further details that support our results.

## Acknowledgments

We thank Brent Doiron for enlightening conversations about this research. We thank the Southern Methodist University (SMU) Center for Research Computing for providing computational resources.

## Author Contributions

C.L., A.K.B., and W.L.S. designed the research. C.L. designed and implemented the computational models. SH. G. and W.L.S. designed and performed the experiments. C.L., A.K.B., SH.G. and W.L.S. drafted the paper. C.L., A.K.B., and W.L.S. edited the paper.

## Input statistics give reasonable model results

For all firing rate models, we fixed the time-varying mean of ORN inputs *μ*(*t*), insuring MC received stronger inputs than PGC and also including odor-evoked increases for GC (Fig. 2**B**, top). GC do not receive direct odor input, but we justify increasing *μ*(*t*) to account for unmodeled excitation from other glomeruli to adequately capture odor-evoked increases in GC firing, but we never allowed the *σ* for GC to be time-varying.

The resulting firing rates depend on specific parameters, but we checked that the set of coupling strengths (Fig. 3**D**) for the models with 3 different sets of noise input resulted in MC spontaneous firing rates being larger than GC and slightly larger than PGC while in the evoked state the MC firing is slightly smaller than PGC but still larger than GC (see Fig. S2**A**). Although our data only has MC recordings, *in vivo* anesthetized recordings of freely breathing rats by the Isaacson lab showed larger spontaneous and evoked firing for MC than GC on average (Cang and Isaacson, 2003). While firing in that study depends on odor concentration among other factors, a visual comparison of their Figures 6 and 10 shows larger evoked firing by MC than GC.

The firing rate relationships in Fig. S2A are consistent with the populations averages in the large and realistic coupled OB model in Li and Cleland (2017), where PGC and MC fire with similar rates that are higher than GC. Furthermore, Wellis and Scott (1990) showed intracellular voltage traces with current injection for both GC and PGC with varying behaviors from cell to cell (like Cang and Isaacson (2003)), but PGC visually appear to fire more spikes (their Figs. 8 and 9) than GC (their Fig. 5), even when GC exhibited strong responses to odor stimulation; they did not report comparisons of firing rates.

## Firing rate model can exhibit evoked decreases in variability

Although the firing rate model we use is relatively simple, we show here that the dynamic range is vast. In particular, circuit parameters could be manipulated to yield a decrease in variability and co-variability with stimulus onset.

Using Monte Carlo simulations, we calculated the steady-state spike count covariance in the spontaneous and evoked states over the entire 10,000 point parameter space of coupling strengths and identified those parameter combinations for which both variance and covariance decreased in the evoked state (Fig. S3**A**, magenta points). We see that the coupling strength values are slightly different for the 3 input noise regimes, with small fixed *σ* having the most such points, following by large fixed *σ*, then time-varying a (Fig. S3i–**iii**). Notice that in all cases, both the independent and common GC inhibition have to be relatively strong while there does not appear to be any dependence on MC excitation to GCs. The actual steady-state values are shown in Figure S3**B**, plotted with the average values (blue squares) and experimental data (black curve).

The model can also exhibit stimulus-evoked decreases in the spike count variance, but to robustly show this in all 3 input noise regimes, we had to consider much larger inhibition strength values: 10 ≤ *w*_*MG*_ 20, 5 ≤ *w*_*Gc*_ 10, and 4 ≤ *w*_*GM*_ 8 (Fig. S3C). There does not appear to be a dependence on the MC excitation (*w*_*GM*_) but there is a dependence on total inhibition (*w*_*MG*_ and *w*_*Gc*_) in that both have to be large (Fig. S3C). In the (*w*_*Gc*_, *w*_*MG*_) plots (Fig. S3**Ci, Cii, Ciii**, lower left sub-panels), we see there is a region of gray points in the upper-right corner corresponding to larger *w*_*Gc*_ and *w*_*MG*_ values where the inhibition is so strong that the mean spike counts do not increase in the evoked state. Notice that in all cases, the common GC inhibition *w*_*Gc*_ spans the range because all that matters for the MC spike variance is the total GC inhibition, in contrast to the covariance where *w*_*Gc*_ has a prominent effect that cannot be counteracted with independent inhibition. Figure S3**D** shows the steady-state spike count variance values in spontaneous and evoked (magenta) along with average values (blue squares) and the experimental data (black curve).

Figure S3 demonstrates that even qualitatively matching our experimental data (i.e., increases in mean and variability at stimulus onset) requires parameter tuning in this model.

## Details of the transfer function calculation

To calculate he transfer functions used in our main firing rate model, we implemented and simulated a large-scale model of OB cells. Our model implementation of all three multi-compartment cell types (MC, PGC, GC) are based entirely on models developed by the Cleland Lab (Li and Cleland, 2013, 2017), with coupling between different cell types mediated by dendrodendritic interactions (Rall et al., 1966). For example, synaptic coupling between MC and GC only occurs at part of the lateral dendrite of MC and part of the dendrite/spine of GC; MC and PGC are coupled only at the MC tuft and PGC dendrite/spine.

The main parameter we vary here and all cells to get the transfer function is the applied current: I_*app*_ (see Eq. (17)–(19)). Each cell model contains different compartments, each having intrinsic ionic currents, including sodium, persistent sodium, potassium delayed rectifier, fast-inactivating A-current, slow-inactivating transient potassium current, L-type calcium current and a calcium-dependent potassium current. All parameters, compartments, intrinsic ionic currents and their gating variables are the same as in (Li and Cleland, 2013, 2017) unless otherwise stated.

## Mitral cell model

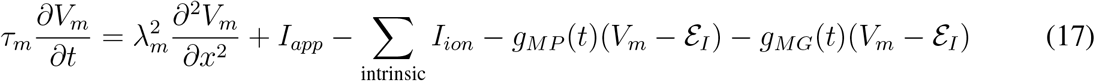

where *x* represents spatial location assumed to be 1-dimensional with no flux boundary conditions, and with four compartments: tuft, apical dendrite, soma, and lateral dendrite.

The complete list of intrinsic ionic currents are: *I*_*Na*_ (sodium), *I*_*NaP*_ (persistent sodium), *I*_*DR*_ potassium delayed rectifier, *I*_*A*_ (fast-inactivating-) and *I*_*KS*_ (slow-inactivating transient potassium current), *I*_*CaL*_ (L-type calcium current), *I*_*KCa*_ (calcium-dependent potassium current). Not all compartments contain all of these ionic currents, see Li and Cleland (2013, 2017) or our code on GitHub.

## Granule cell model

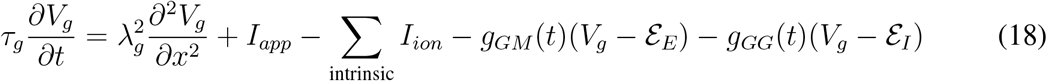

where *x* represents spatial location assumed to be 1-dimensional with no flux boundary conditions, and with two compartments: dendrite/spine, and soma.

## Periglomerular cell model

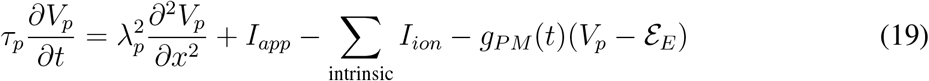

We use standard synaptic connections, with all presynaptic GC and PGC providing GABA_A_ inputs, and all MC providing both AMPA and NMDA – all AMPA and NMDA connection strengths are the same for any given pre/postsynaptic pair, with g_*X*_(*t*)= *w*_*X*_s_*X*_(*t*) where *w*_*X*_ is the strength and all *s*_*X*_ evolve according to:

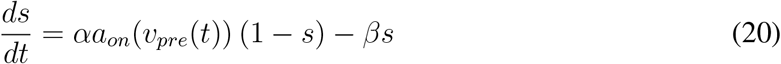

Here *a*_*on*_(*v*_*pre*_) = 1/ (1+ exp (-*v*_*pre*_ - *v*_*t*_)/*v*_*s*_. We adopt the Li and Cleland (2013, 2017) convention of graded inhibitory synapses; GABA_A_ synapses activate via a sigmoidal function with a lower threshold of *v*_*t*_ = −40 mV and scale *v*_*s*_ = 2, while AMPA and NMDA synapes have *v*_*t*_ = 0 mV and *v*_*s*_ = 0.2. The *α* and *β* values are: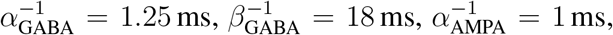, 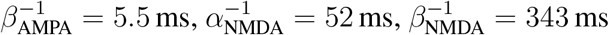.

In Fig. S4**Ai** we see that with an abrupt step of current injection from 0 to 50 *μ*A/cm^2^, the uncoupled PGCs exhibit transient spikes but no sustained spiking, with similar voltage profile as the model in Li and Cleland (2013) (their Fig. 4) and in slice recordings (McQuiston and Katz, 2001). The uncoupled MCs exhibit sub-threshold oscillations as observed in experiments (Desmaisons et al., 1999; Balu et al., 2004), and the number of spikes in a burst increases with current step size (Balu et al., 2004); see Fig. S4**Aii**, again like the model in Li and Cleland (2013). Figure S4**D** shows example voltage trajectories in a coupled network where PGC and GC provide GABA_A_ synaptic inputs and MC provides both AMPA and NMDA inputs with same weight to the appropriate compartments. Voltages within the same cell vary slightly depending on compartment and spatial location. The inset of MC voltage (Fig. S4**D** bottom) shows relatively fast spike propagation, on the order of a few milliseconds (Xiong and Chen, 2002).

## Calculating transfer functions

Firing rate models generally use the transfer functions of individual cell models that are uncoupled. However, since there is no sustained firing in the uncoupled PGC model (Fig. S4**Ai**) with even large values of injected current, the PGC transfer function is not viable. Thus, to incorporate the affects of PGC, we have to instead use transfer functions calculated from reciprocally coupled cells.

We considered two reciprocally coupled networks: i) 1 GC and 1 MC, and ii) 1 PGC and 1 MC; in each network we varied the constant current infection values with fixed synaptic coupling strengths *w*_*GM*_ = *w*_*MG*_ = *w*_*PM*_ = *w*_*MP*_ that are dimensionless; recall *g*_*X*_(*t*) = *w*_*X*_s_*X*_(*t*) where *s*_*X*_ evolves according to Eq. (20) and *g*_*X*_ are in the voltage equations for all 3 cell types. The firing rates were calculated with 50 s of biological time (removing initial 1 s of transients); the average of the two MC firing rates is used.

We calculated several transfer functions (Fig.S4**C**) with fixed synaptic coupling strengths *w*_*GM*_ = *w*_*MG*_ = *w*_*PM*_ = *w*_*MP*_, using values of 60, 80, 100, 120, and 140. Figure S4C shows the resulting curves for each cell type and for the synaptic coupling strengths. Since the input (or constant current injection values) are scaled and the curves appear relatively similar, we use the *w* = 100 curves.

---

**Figure S1:**
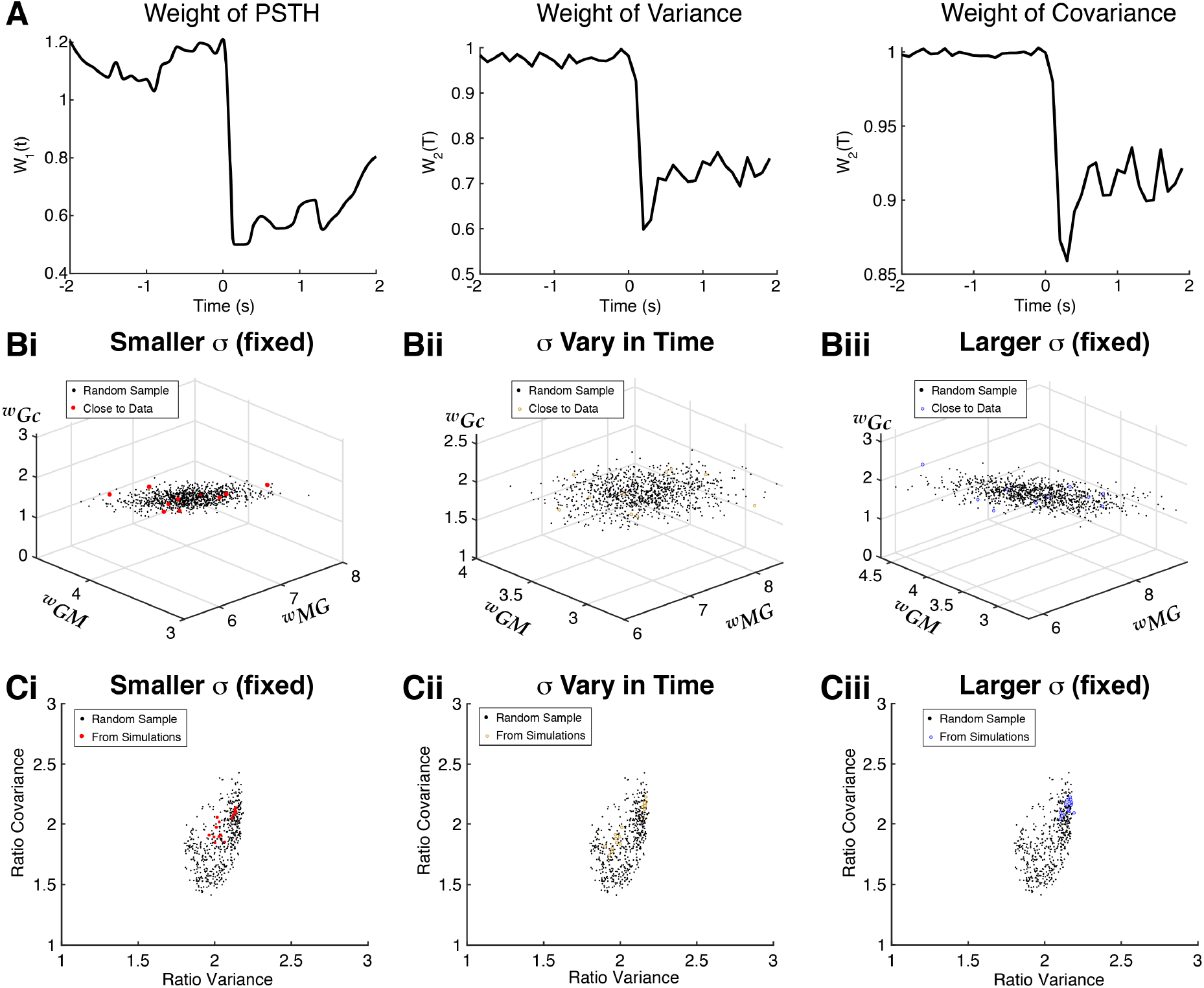
Details for error calculations. **A**) Time-varying weights to calculate error for each of the spike statistics for anesthetized data only (weights are all 1 for awake data). We choose a monotonically decreasing function of the standard deviation of the data (across population / pairs). The weight of the PSTH is *W*_1_(*t*) = 1 + 0.5 tanh ((*std*(*t*) – 10)/5), the spike count variance is *W*_2_(*T*) = 1 + 0.5 tanh ((*std*(*T*) – 1)/5), covariance is: *W*_3_(*T*) = 1 + 0.5 tanh ((*std*(*T*) 0.2)/5). **B**) Black dots are random samples of coupling strength parameters preserving relationships of parameters that best match time-varying statistics of data for the 3 input noise regimes (10 best parameters in Fig. 3**D** are in red, tan, and blue), used for results in Fig. 4. **C**) Scaling factors from simulations (red, tan and blue circles, again from 10 best parameters) relating instantaneous spike count variance and covariance to spike statistics in 100 ms time window. Horizontal axis shows ratio of spike count variance in instantaneous window (dt) to 100 ms time window, vertical shows same ratio for covariance. Each parameter set has 2 ratios: time-averaged ratios in spontaneous state, and time-averaged ratios in the evoked state. We use a single large random sample of ratios (black dots are same for all 3 panels) preserving the dependencies between the variance and covariance ratios; used for results in Fig. 4. Code to generate all random samples on GitHub.

**Figure S2:**
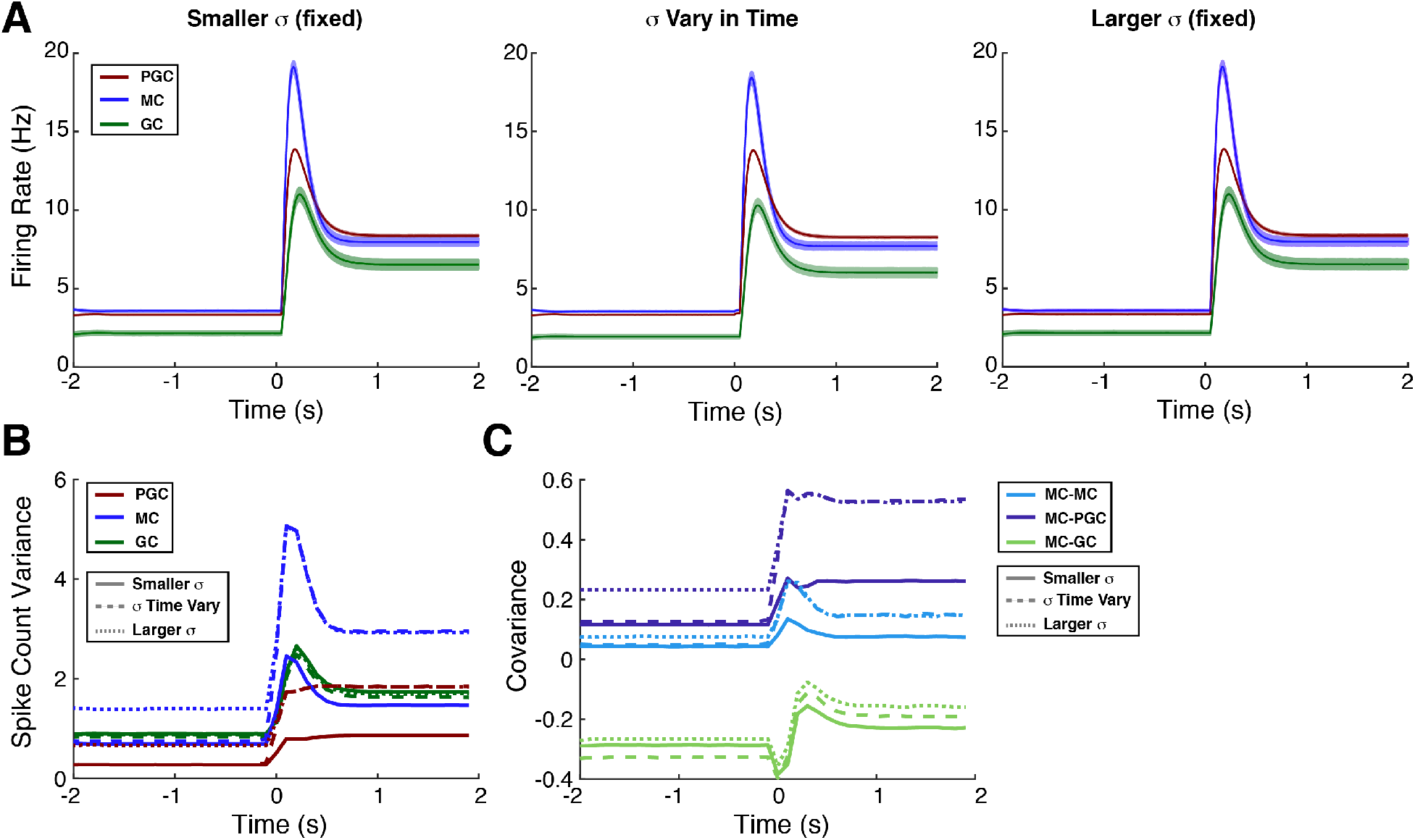
Spike statistics of other cell types in model. Firing rate model results for 3 input noise regimes, averaging over the 10 best sets of coupling strengths (Fig. 3**D** in main text). These model results are consistent with experiments, see **Input statistics give reasonable model results**. **A**) Firing rate for 3 different cell types in the firing rate model, shaded error bars representing standard deviation across 10 best parameter sets. **B**) Spike count variance for 3 different cell types, showing 3 different input noise regimes. **C**) Spike count covariance for 3 pairs of cell types, again combining 3 input noise regimes. Standard deviations (shaded regions) are omitted in **B**) and **C**) because they are small.

**Figure S3:**
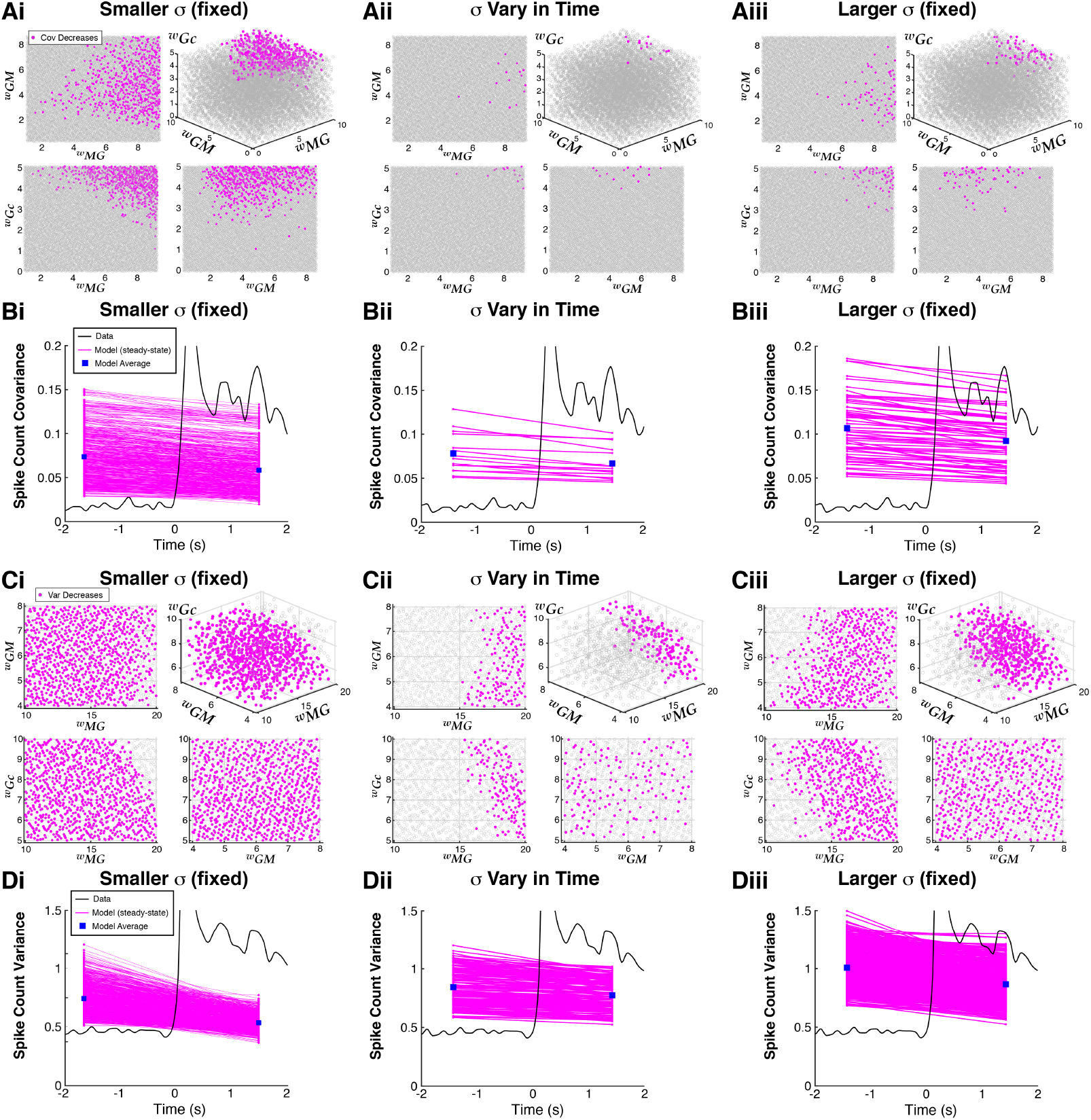
Firing rate model can exhibit odor-evoked decreases in variability. **A**) Parameters where covariance decreases with stimulus and mean spike count increases (magenta dots) from the entire set of 10,000 points and for all 3 input noise regimes. Results vary depending on input noise regime, but both the independent and common GC inhibition must be relatively strong while there does not appear to be a dependence on MC excitation. **B**) Covariance values when evoked is less than spontaneous (magenta), plotted with data (black); the blue squares denote averages of the model results. **C**) With much larger GC inhibition than before, stimulus-induced decreases in variance and increases in mean spiking (magenta dots) can occur. For **C**) and **D**) we test 1000 points with the inhibition values twice as large: 10 ≤ *w*_*MG*_ 20, 5 ≤ *w*_*Gc*_ 10, and a subset of excitation values 4 ≤ *w*_*GM*_ 8. **Ci**) Small fixed *σ* easily results in evoked decreases in variance. For time-varying *σ* (**Cii**) and large fixed *σ* (**Ciii**), the inhibition has to be relatively large for evoked decreases in variance. In all of the (*w*_*MG*_, *w*_*Gc*_) plots, the upper right corner is gray because when total GC inhibition is excessive, the mean spike counts decrease in the evoked state. **D**) All variance values plotted with data (same format as **B**)).

**Figure S4:**
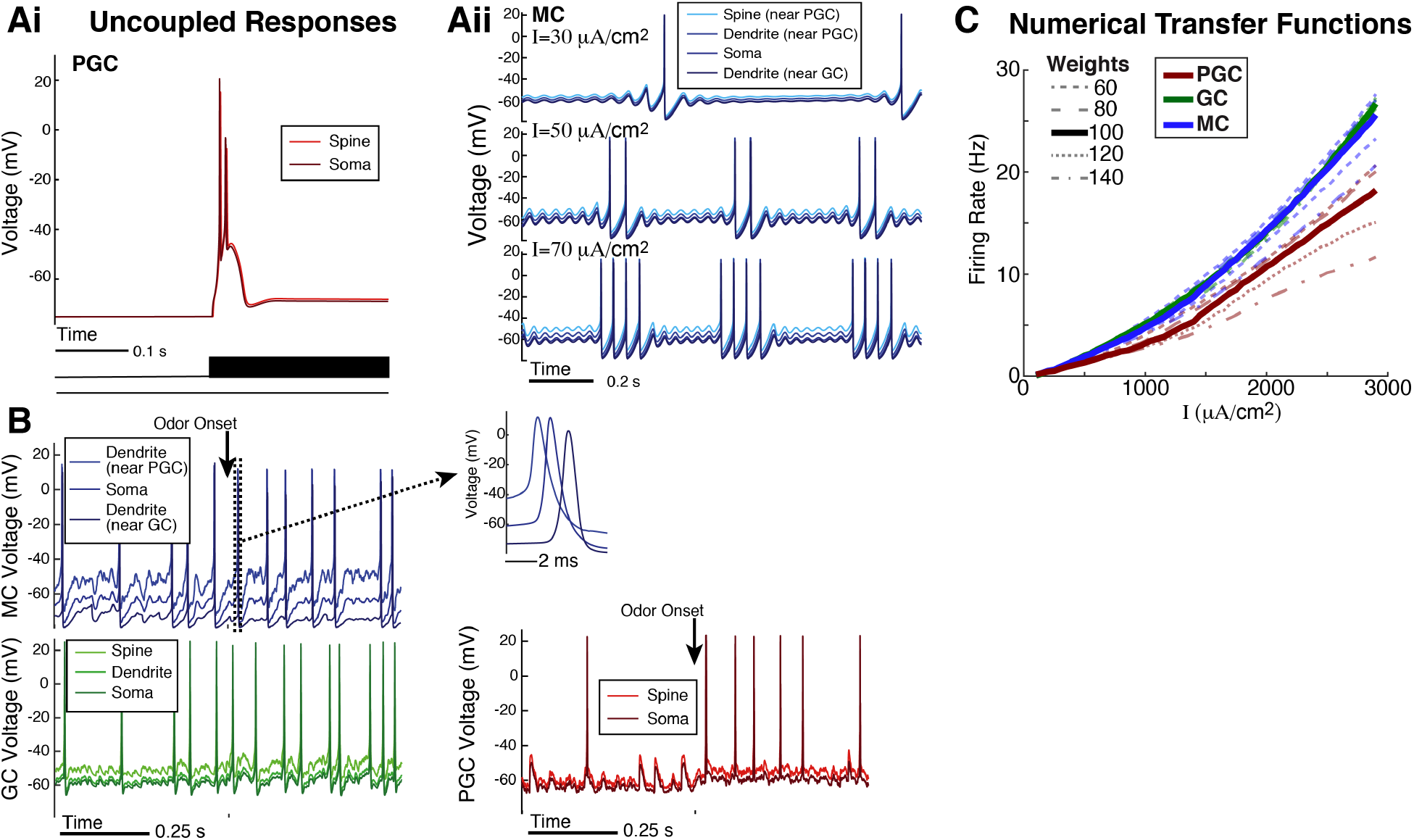
Multi-compartment biophysical OB model used for the firing rate model. Small network of Cleland OB models (Li and Cleland, 2013, 2017). **A**) The firing rate response of uncoupled cells capture some known behavior observed in experiments. i) From rest (no current applied), an abrupt step of current injection to 50 *μ*A/cm^2^ results in transient spikes but no sustained firing, consistent with Li and Cleland (2013); McQuiston and Katz (2001). ii) Uncoupled MC show prominent sub-threshold oscillations (Desmaisons et al., 1999; Balu et al., 2004) and an increase in number of spikes per burst as applied current values increase, consistent with experiments in (Balu et al., 2004). **B**) Voltage traces of the coupled network show random spontaneous and evoked firing; inset of MC at bottom shows relatively fast propagation within a cell, on the order of a few milliseconds (Xiong and Chen, 2002). The results in **A**) and **B**) are not surprising because Li and Cleland (2013, 2017) showed these. **C**) Numerically computed transfer functions with several coupling strengths. Since the shape of the curves are qualitatively similar, we set the curves to be the solid thick curves (*w* = 100) in the firing rate model.

